# The combination of topological data analysis and mathematical modeling improves sleep stage prediction from consumer-grade wearables

**DOI:** 10.1101/2023.10.18.562982

**Authors:** Minki P. Lee, Dae Wook Kim, Olivia Walch, Daniel B. Forger

**Author notes:** These authors contributed equally (Minki P. Lee, Dae Wook Kim).

## Abstract

Wearable devices have become commonplace tools for tracking behavioral and physiological parameters in real-world settings. Nonetheless, the practical utility of these data for clinical and research applications, such as sleep analysis, is hindered by their noisy, large-scale, and multidimensional characteristics. Here, we develop a neural network algorithm that predicts sleep stages by tracking topological features (TFs) of wearable data and model-driven clock proxies reflecting the circadian propensity for sleep. To evaluate its accuracy, we apply it to motion and heart rate data from the Apple Watch worn by subjects undergoing polysomnography (PSG) and compare the predicted sleep stages with the corresponding ground truth PSG records. We find that TFs and clock proxies can improve the overall performance of wake/REM/NREM sleep classification, particularly in identifying REM and NREM sleep (AUROC/AUPRC improvements > 9% and REM/NREM accuracy improvement “ 12%). We find that this improvement is mainly attributed to the heart rate TFs. To confirm this, we compare the heart rate TFs between two groups, expected to have different cardiovascular conditions: younger, healthy subjects from the Apple Watch cohort and elderly subjects from the Multi-ethnic Study of Atherosclerosis cohort. Indeed, TFs largely vary across REM and NREM sleep in younger individuals while the variations disappear in elderly individuals, explaining the enhanced improvements in REM or NREM sleep classification problems only in the younger individuals. This study demonstrates the benefits of combining topological data analysis and mathematical modeling to extract hidden inputs of neural networks from puzzling wearable data.

## Introduction

More than 80% of the population is currently living a shift workstyle, leading to insufficient duration and poor quality of sleep^1^. Notably, this inadequate sleep impacts the quality of life of 70 million individuals in the United States^2^ and thus is considered a growing threat to global public health^3^. This negative effect of sleep disturbance becomes more significant when it arises with chronic diseases such as cancer, diabetes, and mood disorders^1,4,5^. Thus, accurate sleep stage classification, for instance, allowing early diagnosis of sleep disorders, has received attention for more than two decades^6–10^. The gold standard for sleep scoring is polysomnography (PSG), which requires a trained PSG technologist, a sleep lab, and monitoring of multiple physiological parameters^11^. Thus, PSG is not suited for longitudinal and ambulatory sleep tracking in the real word, which can benefit individuals who work in occupations where impairment in alertness is high risk (e.g., nurses and transportation workers). To handle this, wearable devices (e.g., Apple Watch and Fitbit) enabling noninvasive continuous and real-time monitoring of physiological proxies (e.g., rest-activity and heart rate) have been widely exploited^12^. Indeed, with the unprecedented development of machine learning and computing powers, researchers have made meaningful progress in developing algorithms that leverage wearable data to predict sleep stage^8–10,13–15^.

Despite the advances, there is clear evidence that the methods using wearable data have limited accuracy for sleep scoring^9,16–19^. One major reason is that wearable data is highly noisy, nonlinear, and large-scale, resulting in large uncertainty of the sleep stage prediction compared with that using PSG data^20,21^. Indeed, the disparity can be more pronounced in real-world scenarios^22^ and in individuals with sleep disorders^23^.

To tackle the limitation, refining noisy wearable data and extracting the hidden valuable features is necessary. One promising tool for this is topological data analysis (TDA), a data analysis framework exploiting theorems in algebraic topology^24–28^. TDA allows to capture robust topological patterns embedded in a noisy data set^29,30^. Its key idea is that data structure can be characterized by counting holes of different dimensions calculated based on persistent homology theory. This characterized topological information is typically summarized and visualized by the so-called persistence diagram^25,28,29,31,32^. Based on this, several useful topological statistics, such as the maximum persistence of a persistence diagram, have been proposed and applied to analyze the structure of time-series data, such as voices, body motions, and heart rate variability^33–36^. Furthermore, electroencephalogram signals have been analyzed using persistence landscape, which maps persistence diagrams into some functional space, such as Banach space^37,38^. This recent successful application of TDA to clinical time-series data indicates its potential usefulness in analyzing wearable data for sleep scoring.

Another tool for analyzing wearable data and improving sleep scoring is mathematical models based on differential equations. Indeed, several pioneer studies revealed the benefits of mathematical models to identify hidden physiological proxies from wearable data^9,39–45^. Specifically, Walch et al. showed that a clock proxy that describes the circadian propensity for sleep, predicted by the mathematical model of the human circadian clock using wearable data as inputs, increases the accuracy of sleep scoring algorithms^9^. However, its effectiveness with TDA techniques remains to be elucidated.

To study the potential benefits of collaborative use of TDA and mathematical modeling, we developed a neural network sleep scoring framework based on both TDA and mathematical models. To demonstrate its usefulness, we applied it to heart rate and motion data previously collected from the Apple Watch. We found that the topological features and clock proxies from heart rate and motion data collectively improved the classifier’s ability to identify REM and NREM sleep. We also show that this collective improvement can be attributed to the combination of clock proxies and topological features from heart rate data.

Based on this, we further explored the underlying dynamics of heart rate that affect the topological features and illustrate the aging effects on heart rate. Our study illustrates the importance of exploiting TDA and mathematical models together for accurate sleep scoring and demonstrates TDA’s effectiveness in analyzing noisy wearable data.

## Methods

### Dataset description and experimental protocol

Two different datasets were analyzed in this study. First, we used a dataset consisting of Apple Watch data and PSG recordings from participants (mean age = 29.4 ± 8.5 years, range 19-55) recruited by the University of Michigan Sleep and Chronophysiology laboratory. A second dataset consists of publicly available actigraphy and PSG recordings from a population of older participants (mean age = 69.3 ± 8.9 years, range 56-89) recruited as part of the Multi-Ethnic Study of Atherosclerosis (MESA). Below, we describe the specific procedures and experimental protocols for each dataset.

### Apple Watch dataset

39 subjects were recruited by the University of Michigan Sleep and Chronophysiology laboratory for a study aimed at assessing the effectiveness of wearable technologies for sleep analysis^9^. This study was approved by the University of Michigan Institutional Review Board. They excluded individuals with sleep-related, neurological, or psychiatric impairments, and those working night shifts, having recent extensive travel, or experiencing excessive daytime sleepiness. The remaining participants wore Apple Watches (Series 2 and 3) for 7 to 14 days, except during necessary times like charging or showering, collecting heart rate (HR) and motion data. On the last night, participants visited the lab for polysomnography (PSG) recording while simultaneously wearing an Apple Watch transmitting HR and motion data to a server. A sleep medicine physician excluded those with PSG findings suggesting REM sleep disorder or OSA. Data from the remaining 31 participants (average age 29.4 ± 8.5 years, range 19-55) were analyzed. See ^9^ and Supplementary materials for details.

### MESA dataset

6,814 participants consisting of black, white, Hispanic, and Chinese American men and women were recruited from 2000 to 2002 as part of the Multi-Ethnic Study of Atherosclerosis (MESA) for a longitudinal investigation of the factors contributing to the development of subclinical cardiovascular diseases and progression of clinical cardiovascular diseases^46–48^. Among 6,814 recruited participants, 2,237 participants underwent a Sleep Exam from 2010 to 2013, which consisted of 7 days of wrist-worn actigraphy (Actiwatch Spectrum; Philips Respironics, Murrysville, PA) data collection, a sleep questionnaire, and a full overnight unattended PSG recording. Because of the difference in data collection methodology, the HR data was measured using pulse oximetry during PSG recording rather than from actigraphy. Moreover, the MEMS-type actigraphy measured step counts rather than triaxial acceleration. For computational feasibility, PSG recording and actigraphy data from a subpopulation (*n* = 214, mean age = 69.3 ± 8.9 years, range 56-89) of an entire MESA cohort was included for the analysis in this study.

### Feature generation

Three different types of features were derived from both HR and motion data: raw, topological, and clock proxy features. In the following, we explain the methodology and procedure for obtaining each feature.

### Preprocessing and constructing raw HR and motion features

HR data was collected in units of bpm every few seconds (Fig 1A left panel). Because the number of HR measurements within some 30 seconds epoch was sometimes insufficient to perform accurate analysis, cubic spline interpolation was applied to the HR data and resampled it every one second. To obtain the HR feature, resampled HR data was normalized by the 90th percentile in the absolute difference between each measurement and the mean HR over the entire sleep period, as done in previous work^9^. The standard deviation of normalized HR over each 30 second epoch was used as a ‘raw’ HR feature.

**Fig 1.**
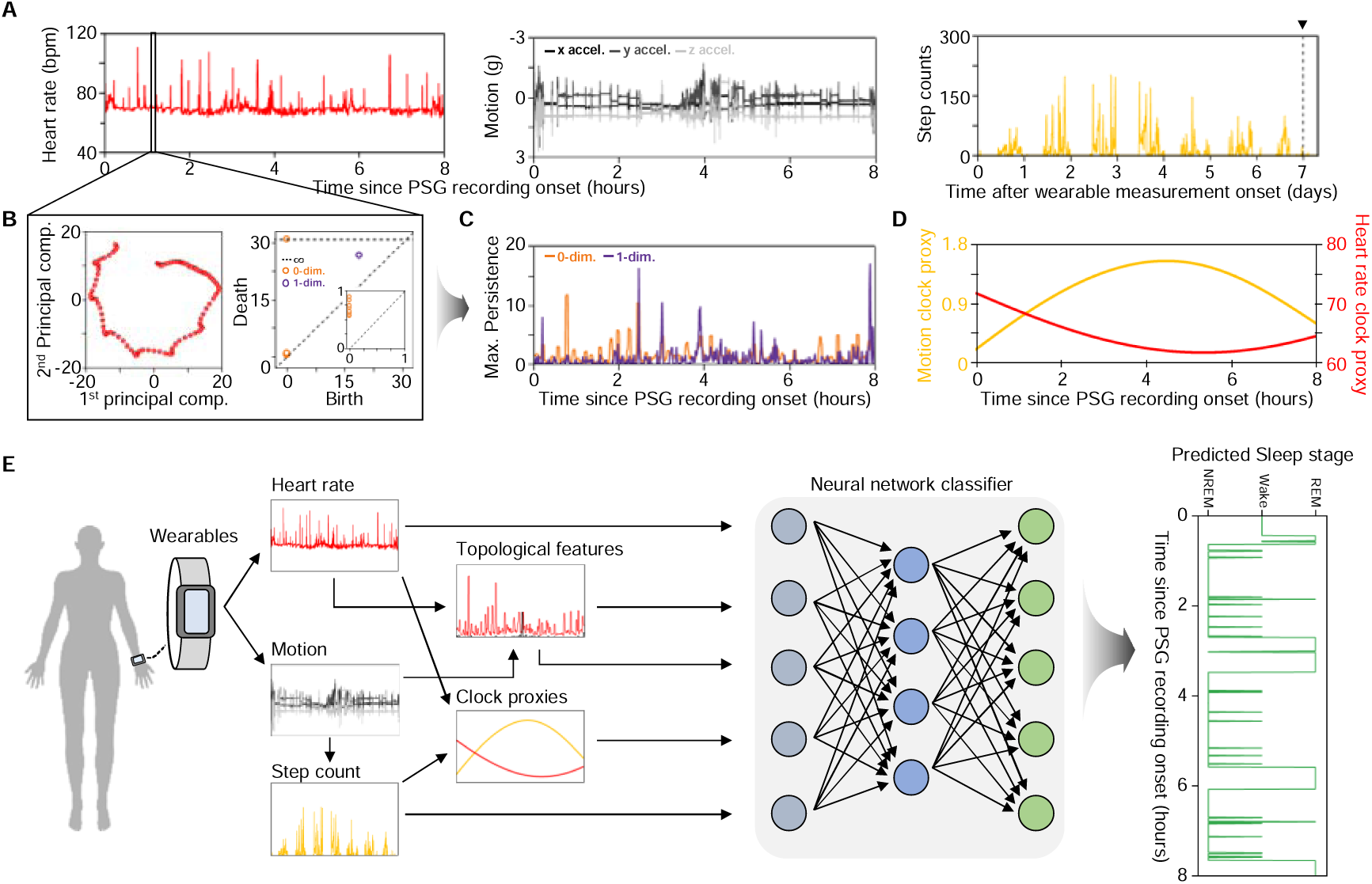
TDA and mathematical modeling extracts hidden proxies from wearable motion and heart rate data to predict sleep stages using a neural network classifier. (A) Participants wore an Apple Watch collecting heart rate (HR) and triaxial acceleration data. At the end of the ambulatory recording period (black triangle at top right), participants showed up to a sleep lab to undergo a PSG recording while simultaneously wearing an Apple Watch. Motion data was used to estimate step counts using a previously developed computational package^49^. **(B)** Wearable HR and motion data were embedded into a higher dimensional space. By selecting appropriate embedding parameters through minimization of the time-delayed mutual information and false nearest neighbors^51^, a point cloud can be constructed from time-series data (left). It is represented by its 1^st^ and 2^nd^ principal components for visualization only. TDA was applied to the embedded point cloud data to construct 0^th^ and 1^st^ order persistence diagram (right). Here, a point further away from the diagonal (e.g., a purple dot) represents a distinct topological feature. **(C)** Maximum 0^th^ and 1^st^ order persistence values from persistence diagram for each epoch were used as topological features (TFs). **(D)** Clock proxies (CPs) were inferred from motion data (yellow) and heart rate data (red) using mathematical models^42,52^. See Methods for details. **(E)** A neural network classifier using raw wearable data, TFs, and CPs as inputs was constructed to predict sleep stages.

Triaxial acceleration motion data from Apple Watch was collected in units of *g* for each *x, y,* and *z* direction (Fig 1A center panel). Cubic spline interpolation was applied to the motion data in each direction, as done in the processing of HR data. The interpolated motion data was then used to estimate the step counts for each 30 second epoch using a previously developed computational package^49^ (Fig 1A right). Finally, the estimated step counts were convolved with a Gaussian (“ = 50 seconds) as done in previous work^9^ (Fig 1A right panel), and convolved step count was used as a ‘raw’ motion feature.

### Implementation of TDA to extract topological features

The central idea behind TDA is to capture robust topological invariants hidden in the data^24–28,31^. Topological invariants can be quantified using the theory of persistent homology, which essentially counts the number of “holes” of different dimensions (i.e., homology) within the point cloud data and records how long they persist. For instance, the 0^th^ order homology refers to the connectivity, 1^st^ order homology refers to the circular hole, and 2^nd^ order homology refers to the void embedded within a collection of data points (see Supplementary materials for details). To capture the persistence of these holes, TDA constructs a ball of growing radius centered at each data point to represent the data as a combinatorial graph (i.e., simplicial complex). As the radius of the balls continuously increases, the simplicial complex continuously evolves, and holes within the simplicial complex appear and disappear (see Supplementary materials for details). The radius at which different holes are born and die is summarized in a so-called persistence diagram^25,28,29,31,32^. Each point on the n^th^ order persistence diagram represents the persistence of distinct n^th^ order holes within the simplicial complex: its *x* coordinate represents the radius value at which n^th^ dimensional holes are born, and the *y* coordinate represents the radius at which holes disappear. Using the persistence diagram, one can visualize topological invariance hidden inside the data to uncover new findings previously unobservable with traditional statistical methods.

We used the persistent homology approach to extract hidden Topological features (TFs) of the Apple Watch HR data. Specifically, for each 30 second epoch, we considered interpolated HR data from 90 second window before and after the 30 second epoch of interest. Together, 210 seconds of time-series HR data was embedded into a higher dimensional space using Takens’s delay embedding map^50^. Here, the optimal embedding dimension d and delay parameter r were determined by minimizing the time-delayed mutual information and false nearest neighbors^51^ of individual subject’s HR data during PSG recording. The embedded HR point-cloud data (Fig 1B left panel) was then used to generate a 0^th^ and 1^st^ order persistence diagram, which captured the hidden topological invariants of the HR data (Fig 1B right panel). The maximum persistence of 0^th^ and 1^st^ order persistence homology, defined as the maximum difference in the *x* and *y* coordinates of all points on the 0^th^ and 1^st^ order persistent diagram (e.g., a purple point on Fig 1B right panel) excluding those on the infinity line, were then used as 0^th^ and 1^st^ order HR TFs, respectively (Fig 1C).

The 0^th^ and 1^st^ order TFs of the HR data are collectively referred to as ‘HR TFs’ throughout this study. Similarly, in the MESA dataset, we computed HR TFs from the HR data measured from pulse oximetry during overnight PSG recording because the participants in the MESA study wore an actigraphy (Actiwatch Spectrum; Philips Respironics, Murrysville, PA), which is incapable of measuring HR data.

Like with the HR data, we obtained 0^th^ and 1^st^ order TFs from the Apple Watch motion data, using Euclidean magnitude of the interpolated triaxial acceleration data to generate a persistence diagram. The 0^th^ and 1^st^ order TFs of the motion data are collectively referred to as ‘motion TFs’ throughout this study.

### Estimation of clock proxies using mathematical models

In addition to the raw features and TFs, we estimated clock proxies (CPs) that approximate the effects of the changing circadian drive to sleep from wearables using mathematical models. We adopted a limit-cycle mathematical model of the human circadian pacemaker^52^ to estimate circadian signals from wearable data, as in previous work^9^. The model requires light measurement (in lux) as an input. However, the current generations of consumer-grade wearable devices do not provide light sensors. To circumvent this, we alternatively used the preprocessed step counts from motion data to impute light measurement, as done in previous work^9,43^. Specifically, we adopted a simple piecewise function that converts step counts into different light levels depending on the timing of the day: if a measured step count exceeded a user-specified threshold, the estimated light exposure was 50 lux during the night (i.e., 10pm to 7am), 500 lux during the morning and evening (i.e., 7am to 10am and 4pm to 10pm), and 1000 lux during the day (i.e., 10am to 4pm). If a step count was below the threshold, it was assumed that the light exposure was 0 lux. In this study, we used 20 steps per minute as a threshold. The converted light input was used to simulate the mathematical model, which outputs a ∼24hr oscillatory trajectory that estimates a circadian propensity for sleep (yellow curve, Fig 1D). The output from the model was used as a proxy for circadian signal, referred to as a ‘motion CP’.

In addition, we predicted a circadian signal in HR from the wearable activity and HR data using a recently developed well-validated mathematical model of the human circadian rhythm in HR^42,45,53,54^. The model describes a basal HR, ∼24hr harmonic oscillation of circadian rhythms in the HR, the effect of activity on the HR, and external factors such as hormones^55^, stress^56^, or caffeine^57^, that may further affect the HR. Specifically, HR at time *t* is described by a harmonic regression and an autocorrelated noise process:

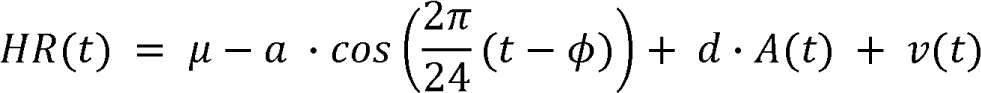

where

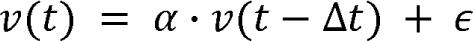

µ represents the basal HR; a represents the circadian amplitude; 4> denotes the timing of the circadian HR minimum; d represents the increase in HR per step (i.e., HRpS); v(t) is a first order autocorrelated noise process that carries over a fraction a of the noise at time t-Δt; and € is a Gaussian random variable with a zero mean. The model was fitted to the measured HR and step counts using the Goodman and Weare affine invariant Markov chain Monte Carlo (MCMC) method^42,58^, and the best estimates of parameters were obtained. Using the estimated parameters, the harmonic component of the model was used as a proxy for the circadian signal, referred to as an ‘HR CP’ (red curve, Fig 1D).

### Neural network learning and performance metrics

A neural network model was used to classify the sleep stage of each 30 second epoch as either (i) Sleep or Wake and (ii) Wake or Rapid Eye Movement (REM) or non-REM (NREM) sleep, using different features (Fig 1E and Supplementary Table S1). The model consists of three hidden layers with an inverse tangent activation function and an output layer with a sigmoid activation function. All hyperparameters of the neural network classifier were obtained by maximizing the overall Area under the Receiver Operating Characteristic curve (AUROC) and the Area under the Precision and Recall curve (AUPRC) (see Supplementary materials for details). The same hyperparameters were used in all neural network training and validations. To evaluate the performance of the neural network model, we used AUROC and AUPRC as primary metrics. Other metrics, including the overall accuracy, F1 score, and Cohen’s K, were further used to analyze classification performance (see Supplementary materials for details).

### Examining the contributions of the features

To systematically investigate how different features contribute to the classification performance, we considered three different realistic scenarios based on the availability of motion and HR data. In the first case, we consider a scenario in which only motion data is available, and only raw motion, motion TFs, and a motion CP were used to represent each epoch. In the second scenario, only HR data is available, and we used raw HR, HR TFs, and a HR CP. Because the motion data is not available, we assumed that HRpS equals to zero in our computation of a HR CP. Finally, we consider a third scenario in which both motion and HR data are available, and all features were used for the sleep stage classification.

### Validation against PSG

We implemented Monte Carlo Cross Validation (MCCV) to generate training and testing data for sleep/wake classification within the Apple Watch dataset. In all scenarios, 70% of all 30-second epochs across the entire Apple Watch dataset were randomly selected as a training set, and remaining 30% of the data were used as a testing set. To account for the randomness in splitting the training and testing dataset, this process was repeated 50 times, and model outputs from all 50 iterations were aggregated to evaluate performance metrics. In all repetitions, no data point was ever used for both training and testing.

We repeated the same procedure to investigate how the TFs and CPs contribute to a more detailed three stages wake/REM/NREM classification. We randomly selected 70% of all epochs from the Apple Watch dataset as a training set and the remaining 30% as a testing set across 20 independent iterations. To analyze the classifier’s performance in wake/REM/NREM differentiation, one-versus-rest ROC and PRC curves were constructed by selecting a positive class and treating other sleep stages as negative classes. This process was repeated for all three sleep stages to investigate the classifier’s ability to distinguish a specific stage (i.e., a positive class) from other stages (i.e., negative classes).

To further validate the combined contribution of TFs and CPs to wake/REM/NREM differentiation, we evaluated the REM/NREM accuracy, overall accuracy, F1 score, and Cohen’s K. Evaluating these metrics requires a careful choice of thresholds to distinguish each epoch based on their computed probabilities. For this, following a method previously employed in ^9^, we first chose a threshold that achieves a desired false positive rate of wake. Then, based on the first threshold, we chose a second threshold to classify all epochs that were not categorized as wake, with the aim of minimizing the difference between the true positive rate for REM and NREM. This process was repeated across a full range of possible false positive rates of wake (i.e., from 0 to 1) at an increment of 0.025. This enables the reduction of degrees of freedom in possible combinations of thresholds so that the performance improvements from each feature can be captured more clearly. Furthermore, this method for threshold selection can account for class imbalance, which typically does not result in the highest overall accuracy.

We implemented Leave-one-subject-out-cross-validation (LOSOCV) to investigate subject-by-subject variability in classification performance. Specifically, for a given subject, we used the data from all remaining subjects to train a classifier and evaluated it on an excluded subject. To prevent overfitting due to class imbalance, 70% of all data from the remaining subjects were randomly used to train a classifier. This process was repeated 20 times for each subject to account for the randomness. For each subject, the output from all 20 iterations was used to compute the same set of metrics, where thresholds were chosen to fix wake accuracy at 60%.

### Local periodicity quantification

We quantified the periodicity of HR data for each epoch using a state-of-the-art periodicity quantification method called Sliding Windows 1-dimensional Persistence (SW1PerS), which successfully detected periodicity from different biological data, such as rhythmic gene expression data, in shape-agnostic manners^30,59^. SW1PerS quantifies the periodicity of the time-series data in a score from 0 to 1, where 0 corresponds to complete aperiodicity, and 1 corresponds to perfect periodicity.

## Results

### Sleep/Wake classification

The neural network model was first trained and tested for sleep/wake classification using the features extracted from wearable data (see Methods for details and Fig 1). ROC and PRC curves and their AUC values summarizing the classifier’s performance are shown in Supplementary Fig S1 and Table S2. When using a raw motion feature alone for training and testing, the model achieved an AUROC and AUPRC value of 0.810 and 0.420, respectively. When using raw motion and CP features together, the AUROC and AUPRC values were improved by 6% and 15% compared to the baseline performance using only a raw motion feature. However, adding in motion TFs did not contribute to differentiating sleep and wake. The results suggest that a motion CP is important for differentiating sleep and wake.

Next, only features derived from HR data were used to train and test the classifier. Unlike motion TFs, HR TFs successfully improved sleep/wake classification. Moreover, the combined use of HR TFs and a HR CP further improved classification. Despite the performance improvements from using HR TFs, the impact of TFs was diminished when exploiting both motion and HR data. On the other hand, CPs continued to effectively enhance performance. These results indicate that CPs mainly contributed to the improvements in sleep/wake classification, while TFs made limited contributions to sleep/wake classification when both motion-based and HR-based features were used.

### Wake/REM/NREM classification

We performed the same procedure to examine the extent to which TFs and CPs contribute to Wake/REM/NREM classification (see Methods for details). To illustrate the classifier’s performance, one-versus-rest ROC and PRC curves and their AUC values are shown in Figs 2 and 3 and Table 1. When only motion-based features were used, we observed similar contributions of motion TFs and a motion CP as in sleep/wake classification: a motion CP plays a critical role in improving the classifier’s performance across sleep stages while motion TFs made a minimal contribution (Fig 2A-C and Fig 3A-C).

**Fig 2.**
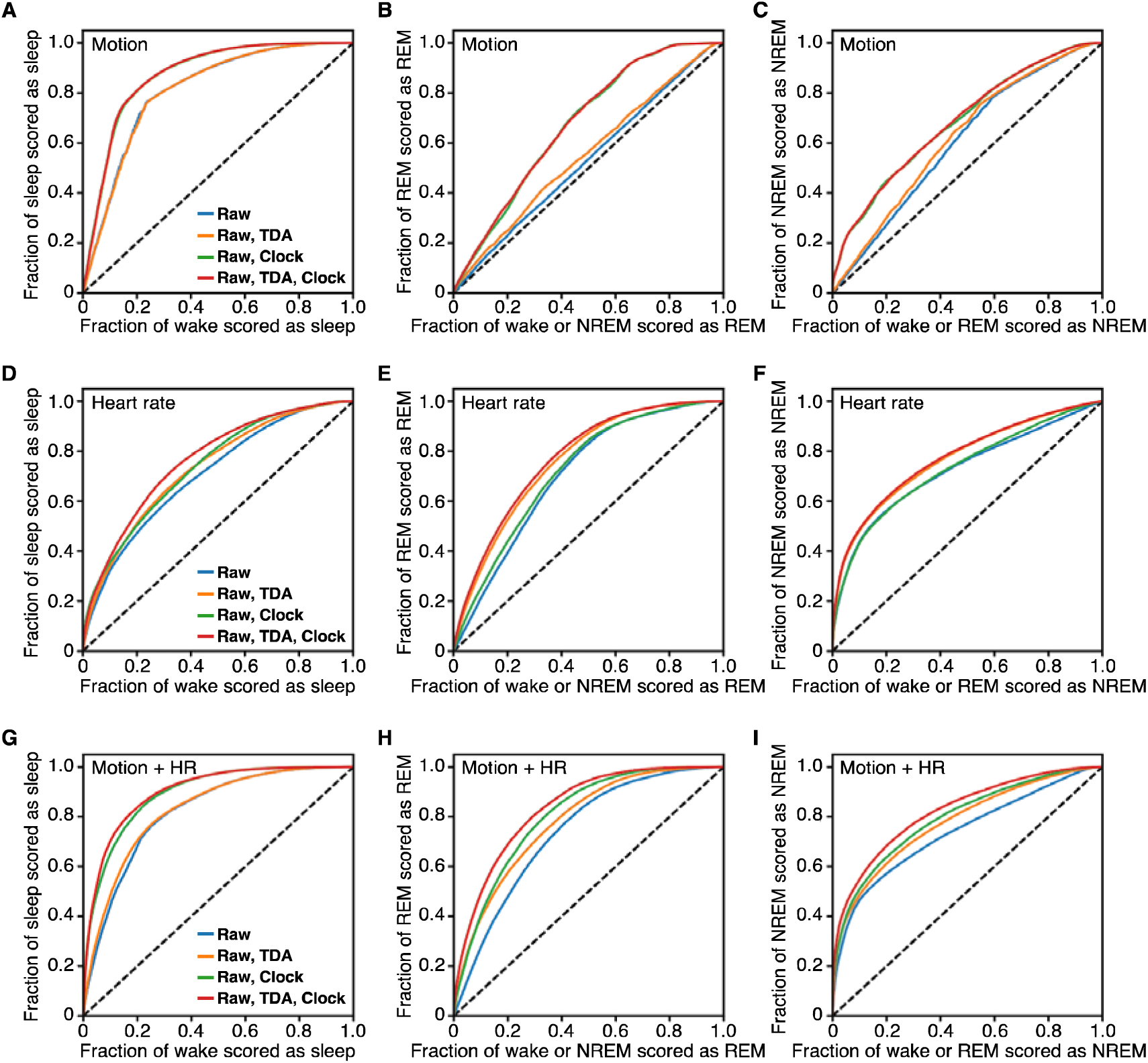
One-versus-rest ROC curves for Wake/REM/NREM differentiation using different features based on the availability of motion and HR data. (A-C) One versus rest ROC curves were generated when only motion data is available. AUROC values for all three stages increased after adding in motion CP, while motion TFs made minimal contributions to differentiating sleep stages. **(D-F)** One versus rest ROC curves when only HR data is available. HR TFs improved overall classifier’s performance, and HR CP mainly contributed to distinguishing wake versus sleep. **(G-I)** One versus rest ROC curves using both motion and HR data. TFs improved REM and NREM AUROC values, and their combination with CPs resulted in further improvements in overall classifier’s performance. The AUROC values are shown in Table 1.

**Fig 3.**
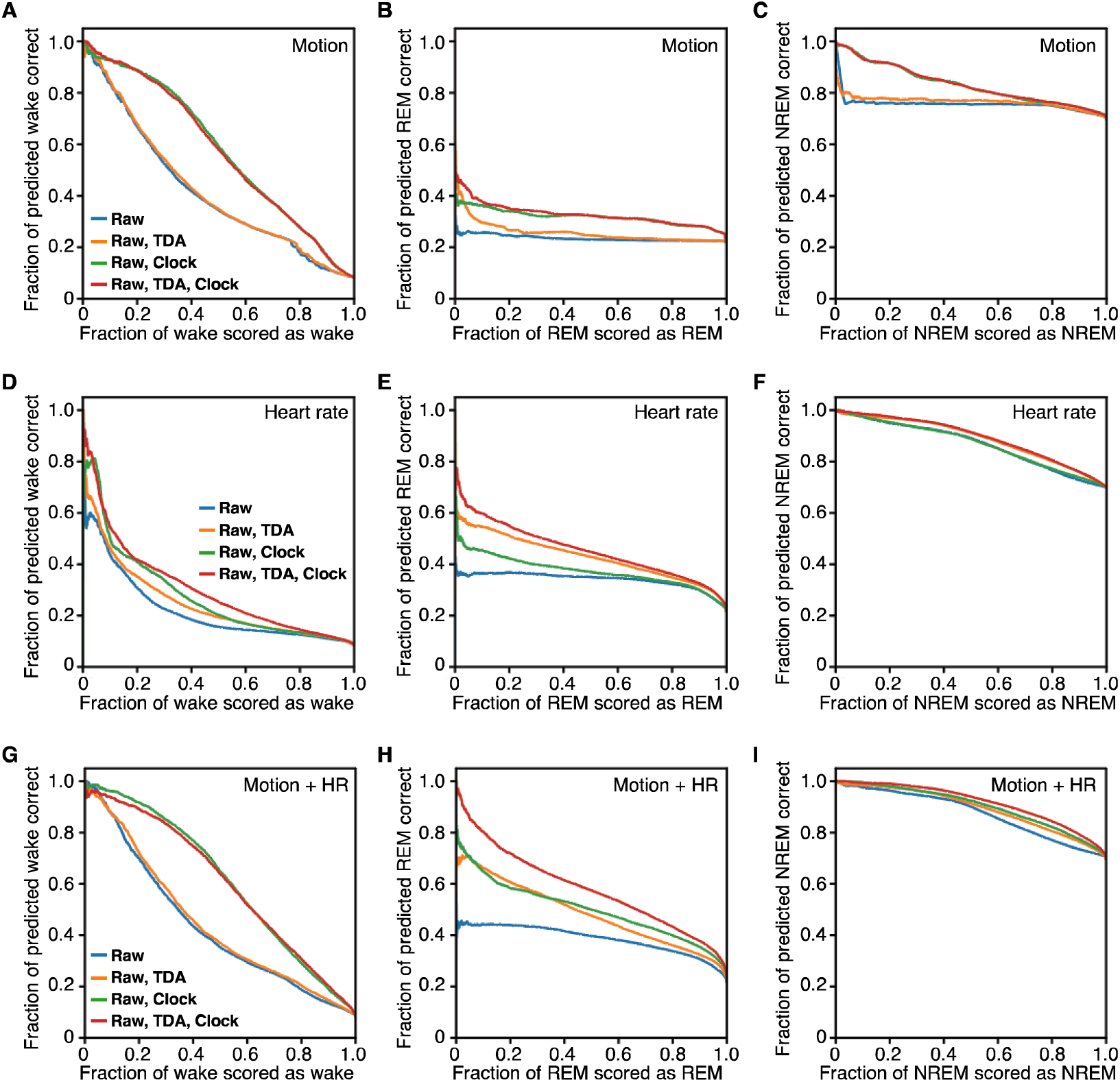
One versus rest PRC curves for Wake/REM/NREM differentiation using different features based on the availability of motion and HR data. (A-C) One versus rest PRC curves were generated when only motion data is available. As with ROC curves, AUPRC values for all three stages increased after adding in motion CP, while motion TFs made minimal contributions. **(D-F)** One versus rest PRC curves when only HR data is available. HR TFs and HR CP mostly contributed to identifying REM sleep. **(G-I)** One versus rest PRC curves using both motion and HR data. The combination of TFs and CPs improved Wake/REM/NREM classification, especially in differentiating REM and NREM sleep. The AUPRC values are shown in Table 1.

**Table 1.** Wake/REM/NREM sleep prediction results within the Apple Watch dataset. The reported AUROC and AUPRC values were calculated from the one-versus-rest ROC and PRC curves for Wake/REM/NREM differentiation within the Apple Watch dataset using the MCCV (Figs 2 and 3).

|  | Types of features used |  |  |  |
| --- | --- | --- | --- | --- |
|  | Raw | Raw, TDA | Raw, clock | Raw, TDA, clock |
| **Motion only** |  |  |  |  |
| **Wake vs not wake** |  |  |  |  |
| AUROC | 0.808 | 0.814 | 0.873 | 0.873 |
| AUPRC | 0.417 | 0.424 | 0.578 | 0.575 |
| **REM vs not REM** |  |  |  |  |
| AUROC | 0.532 | 0.554 | 0.674 | 0.678 |
| AUPRC | 0.236 | 0.258 | 0.318 | 0.327 |
| **NREM vs not NREM** |  |  |  |  |
| AUROC | 0.606 | 0.622 | 0.683 | 0.686 |
| AUPRC | 0.756 | 0.766 | 0.831 | 0.832 |
| **HR only** |  |  |  |  |
| **Wake vs not wake** |  |  |  |  |
| AUROC | 0.714 | 0.742 | 0.750 | 0.777 |
| AUPRC | 0.220 | 0.249 | 0.276 | 0.304 |
| **REM vs not REM** |  |  |  |  |
| AUROC | 0.705 | 0.760 | 0.721 | 0.772 |
| AUPRC | 0.340 | 0.428 | 0.374 | 0.455 |
| **NREM vs not NREM** |  |  |  |  |
| AUROC | 0.723 | 0.768 | 0.730 | 0.774 |
| AUPRC | 0.867 | 0.888 | 0.868 | 0.892 |
| **Motion + HR** |  |  |  |  |
| **Wake vs not wake** |  |  |  |  |
| AUROC | 0.827 | 0.838 | 0.894 | 0.899 |
| AUPRC | 0.433 | 0.446 | 0.611 | 0.603 |
| **REM vs not REM** |  |  |  |  |
| AUROC | 0.738 | 0.782 | 0.806 | 0.837 |
| AUPRC | 0.386 | 0.483 | 0.500 | 0.580 |
| **NREM vs not NREM** |  |  |  |  |
| AUROC | 0.737 | 0.778 | 0.795 | 0.823 |
| AUPRC | 0.875 | 0.894 | 0.902 | 0.917 |

Unlike motion TFs, HR TFs made significant improvements to the wake/REM/NREM classification. Specifically, HR TFs improved the one-versus-rest AUROC value by 3.8%, 5.5%, and 4.5% when wake, REM, and NREM sleep were considered as a positive class, respectively (orange vs blue curves, Fig 2D-F and Table 1). Likewise, the AUPRC value was increased by 2.9%, 8.8%, and 2.1% with wake, REM, and NREM as a positive class, respectively (orange vs blue curves, Fig 3D-F and Table 1). In contrast, a HR CP had less significant contributions, particularly when identifying REM and NREM sleep from other stages (e.g., ∼1.5% improvements in AUROC and AUPRC values). These findings indicate that HR TFs are a key factor in improving the classifier’s performance across sleep stages.

In general, combining motion-based and HR-based features further improved wake/REM/NREM classification across all stages (Figs 2G-F and 3G-F and Table 1). Specifically, when identifying REM sleep using just raw motion and HR features, the classifier achieved an AUROC of 0.738 and an AUPRC of 0.386, respectively (blue curve; Figs 2H and 3H and Table 1). When both TFs and CPs were used as additional input features for the classifier, there was a notable performance improvement, with ∼10% increase in AUROC and ∼20% increase in AUPRC (red curve, Figs 2H and 3H and Table 1). These improvements are particularly noteworthy because they surpass the improvements achieved when either TFs or CPs were used as inputs alone (orange and green curves, Figs 2H and 3H and Table 1). A similar trend was observed when identifying NREM sleep: combining of TFs and CPs synergistically improved the classifier’s performance from its baseline performance (Figs 2I and 3I, and Table 1). These synergistic improvements appear primarily attributed to HR TFs and a motion CP, especially considering the limited contributions of motion TFs and a HR CP (Figs 2A-F and 3A-F). These results indicate that extracting latent input features through TDA and mathematical modeling is essential for harnessing the full potential of noisy wearable data collected in real-world settings, particularly in the field of sleep analysis.

### Further validation using different performance metrics

We further confirmed our findings by using additional ROC curves replacing true positive rates with accuracy values that approximately equate the performance between REM and NREM sleep (see Methods for details and ^9^). This summarizes the performance in all three classes. We found that the combination of TFs and CPs improved the classifier’s ability to identify REM and NREM sleep across all possible choices of thresholds (Fig 4A). In particular, when the accuracy of wake was fixed at 60%, TFs and CPs improved the average REM/NREM accuracy by 3% and 9%, respectively, compared to using just raw features (Fig 4A black triangle and Table 2). Note that the improvement from CPs was greater than using TFs because TFs mainly differentiated just REM and NREM sleep, while CPs also improved identification of wake (Fig 2G-I and 3G-I). Combining TFs and CPs improved REM/NREM accuracy by ∼12% compared to the baseline performance (Table 2). Likewise, we evaluated the overall accuracy, F1 score, and Cohen’s K statistics for all possible choice of thresholds and observed similar trends in improvements (Fig 4B-D). Notably, TFs and CPs collectively improved Cohen’s K by ∼14% when the accuracy of the wake was fixed at 60% (Fig 4D black triangle and Table 2).

**Fig 4.**
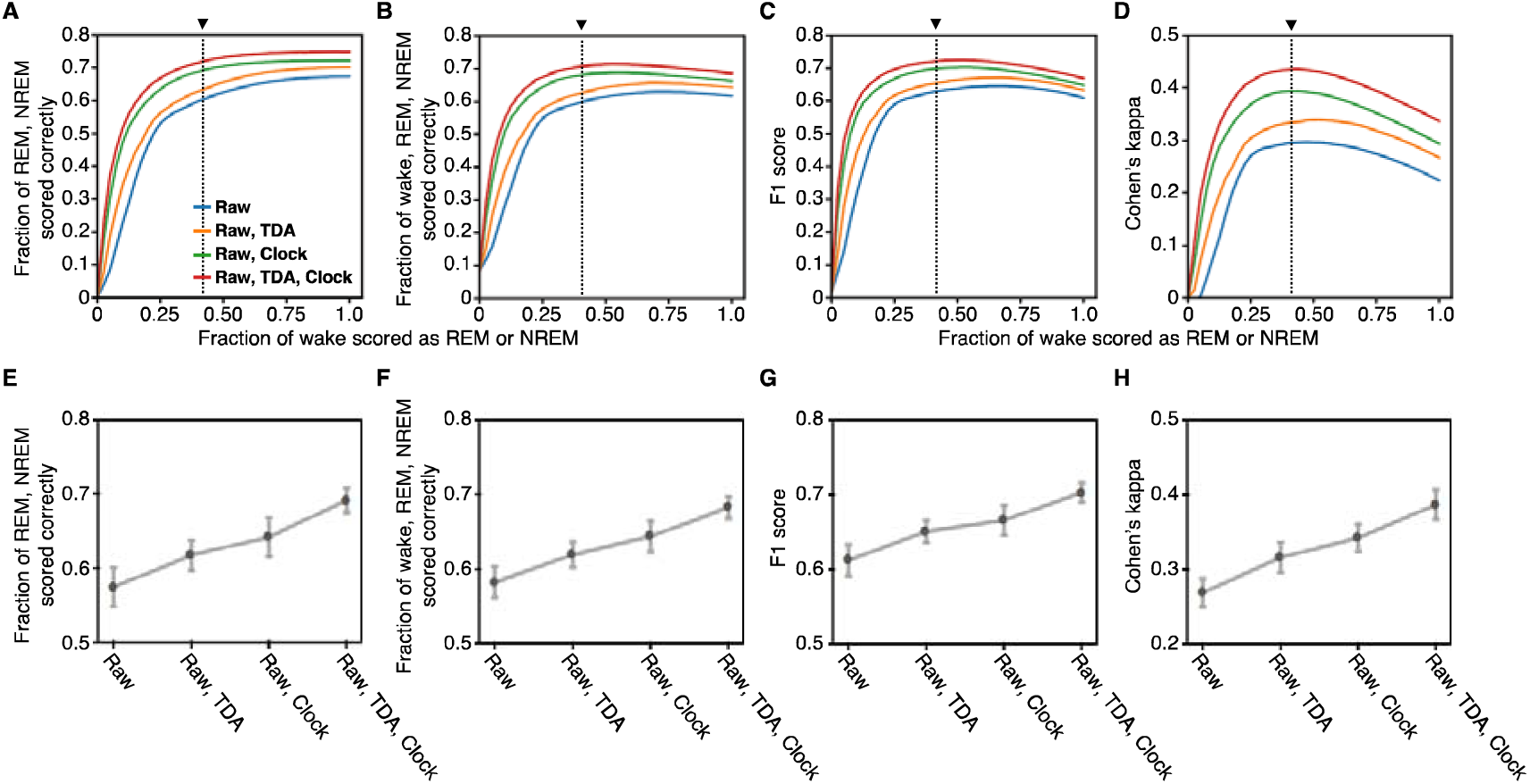
Performance quantification using different metrics. (A-D) REM/NREM accuracy (A), overall accuracy (B), F1 score (C), and Cohen’s K (D) summarizing the performance in all three sleep stage. A binary search was performed to find a threshold that fixes a desired accuracy of wake and another threshold that minimizes the difference between REM and NREM accuracy. Using selected thresholds across full range of desired wake accuracy, average REM/NREM accuracy (A), overall accuracy (B), F1 score (C), and Cohen’s K (D) were evaluated using both motion and HR data. Topological features and clock proxies synergistically improved all performance metrics. **(E-H)** LOSOCV was performed to investigate the inter-individual variability in classifier’s performance. The mean and standard error of the mean (SEM) of REM/NREM accuracy (E), overall accuracy (F), F1 score (G), and Cohen’s K (H), calculated across all validations for each subject, are reported. Here, the error bars represent the SEM.

**Table 2.** Evaluation of different performance metrics of Wake/REM/NREM classification within the Apple Watch dataset. The reported performance metrics were calculated using thresholds that fix the wake accuracy at 60% and minimize the difference between REM and NREM accuracies (Fig 4A-D). The best overall accuracy, best F1 score, and best Cohen’s K from a full range of thresholds search are also reported.

|  | Types of features used |  |  |  |
| --- | --- | --- | --- | --- |
|  | Raw | Raw, TDA | Raw, clock | Raw, TDA, clock |
| Wake correct | 0.6 | 0.6 | 0.6 | 0.6 |
| REM correct | 0.598 | 0.628 | 0.689 | 0.716 |
| NREM correct | 0.598 | 0.627 | 0.688 | 0.716 |
| Overall accuracy | 0.598 | 0.625 | 0.681 | 0.706 |
| F1 score | 0.626 | 0.651 | 0.697 | 0.720 |
| Cohen's $\kappa$ | 0.295 | 0.333 | 0.393 | 0.434 |
| Best accuracy | 0.629 | 0.657 | 0.689 | 0.713 |
| Best F1 score | 0.645 | 0.671 | 0.702 | 0.725 |
| Best Cohen's $\kappa$ | 0.297 | 0.339 | 0.393 | 0.435 |

Next, we employed LOSOCV to investigate subject-by-subject variability in wake/REM/NREM classification performance. The variability in performance using different features is shown in Figure 4E-H and Table 3. On average, REM/NREM accuracy and overall accuracy were improved by ∼4% using TFs and ∼6% using CPs across all subjects (Fig 4E,F and Table 3). These amounted to a total of ∼11% improvement when exploiting all features, compared to the baseline performance. A similar trend in improvements was observed for F1 score and Cohen’s K (Fig 4G,H and Table 3). Notably, Cohen’s K was improved by ∼12% across different subjects when using both TFs and CPs. These results validate that the combination of TDA and mathematical modeling can improve sleep stage prediction even across different individuals.

**Table 3.** Subject-by-subject variation in Wake/REM/NREM classification results within the Apple Watch dataset. The reported performance metrics were calculated using thresholds that fix the wake accuracy at 60% and minimize the difference between REM and NREM accuracies (Fig 4E-H). The results are reported in the form “Mean±SEM”, where the mean and standard error of the mean were calculated across all validations for each subject.

|  | Types of features used |  |  |  |
| --- | --- | --- | --- | --- |
|  | Raw | Raw, TDA | Raw, clock | Raw, TDA, clock |
| Wake correct | 0.6 | 0.6 | 0.6 | 0.6 |
| REM/NREM correct | 0.575±0.026 | 0.618±0.020 | 0.643±0.026 | 0.692±0.016 |
| Overall accuracy | 0.583±0.021 | 0.620±0.017 | 0.644±0.020 | 0.683±0.014 |
| F1 score | 0.613±0.021 | 0.651±0.015 | 0.666±0.020 | 0.703±0.013 |
| Cohen's $\kappa$ | 0.270±0.019 | 0.317±0.020 | 0.343±0.018 | 0.387±0.020 |

### Topological features of heart rate across sleep stages

To gain insight into the contribution of TFs to the differentiation of sleep stages, especially REM and NREM sleep, we investigated how TFs were distributed for each sleep stage. This analysis was first conducted using the Apple Watch dataset, which includes younger and healthy subjects. We estimated the probability density distribution of 0^th^ and 1^st^ order maximum persistence values, referred to as TFs earlier, across sleep stages using Gaussian kernel density estimation (Fig 5). The distributions of motion TFs across different sleep stages were not distinguishable (Fig 5A-C), which matches our results that motion TFs did not contribute to sleep/wake or wake/REM/NREM classification. In contrast, the distributions of HR TFs showed clear differences between REM and NREM sleep (Fig 5D-F). This explains why HR TFs enhanced the performance in predicting REM and NREM sleep.

**Fig 5.**
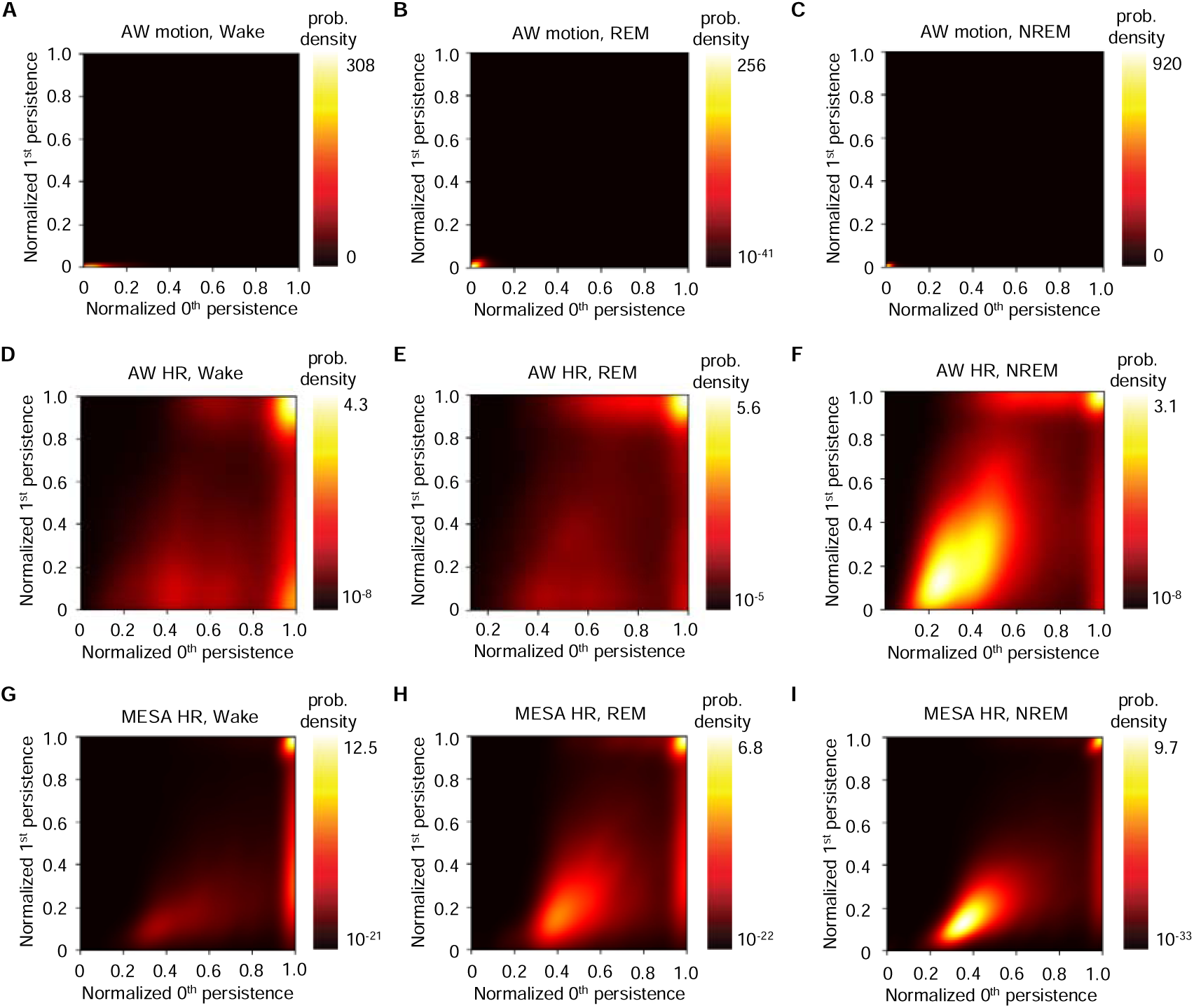
Probability density distributions of TFs. (A-C) Smooth kernel density estimate of probability distributions of motion TFs for each sleep stage in the Apple Watch (AW) dataset. The distributions of motion TFs were indistinguishable across different stages. **(D-F)** Probability distributions of HR TFs in the AW dataset were computed for each sleep stage. The distribution of HR TFs for REM (E) and NREM (F) are distinguishable. **(G-I)** Same process was repeated with the MESA HR data. Unlike the AW dataset, the distribution of MESA HR TFs for REM (H) and NREM (I) sleep are harder to differentiate. Here, 0^th^ and 1^st^ order persistence values were normalized by dividing with 90^th^ percentile value in 0^th^ and 1^st^ order persistence, respectively^9^.

We next explored the distribution of HR TFs in the MESA dataset. The MESA dataset includes a much older population (age (mean±SD) = 69.3 ± 8.9 years, range 56-89) compared to the Apple Watch dataset (age (mean±SD) = 29.42 ± 8.52 years, range 19-55).

It is designed to explore factors related to the development of subclinical cardiovascular disease and the progression to clinical cardiovascular disease. Thus, it is expected that there would be differences in cardiovascular conditions between the two cohorts. This provides an opportunity to study the role of topological dynamics in HR across sleep stages in distinct situations. Indeed, unlike the Apple Watch HR data, the density estimates of HR TFs from the MESA HR data showed no noticeable differences across sleep stages (Fig 5G-I). This matches the negative effect of aging on HR rhythmicity (see ^60–62^ and below). We further examined the HR TFs in relation to demographics, sex, and race (Supplementary Fig S2), because the differences across sleep stages can be obscured by demographic variables^63,64^. We observed no significant differences between the distributions of HR TFs across different subgroups based on sex or race (Supplementary Fig S3 and S4). Consistent with the findings, we noticed no noticeable improvements in wake/REM/NREM classification from using TFs when we trained a neural network classifier from the Apple Watch dataset and validated it on the MESA dataset (Supplementary Fig S5). This indicates that TDA can be used to uncover hidden differences in physiological signals among populations with different backgrounds (e.g., age and medical conditions).

To further understand the differences in TDA’s ability to distinguish sleep stages in younger and older populations, we investigated how the dynamics underlying the HR data differed between the Apple Watch and MESA datasets. Specifically, we calculated the local periodicity and the maximum differences in the HR data for each epoch in both datasets because they are key determinants of the circularity and size of the high dimensional holes generated by the delay embedding, respectively, and therefore the value of TFs (See ^30,59^, Methods and Supplementary Fig S6 for details). Overall, the local periodicity score of the Apple Watch HR data was significantly higher than the MESA HR data (*p*-value < 10^-3^, two-tailed T-test; Fig 6A). Importantly, HR data during REM and NREM sleep measured from the Apple Watch subject was significantly more periodic than the HR data during REM and NREM sleep from MESA subjects (*p*-value < 10^-3^, two-tailed T-test; Fig 6B and C). We next quantified the maximum differences in HR data for each epoch. As illustrated in Fig 6E, the probability distribution of the maximum difference in HR for REM and NREM was clearly different in the Apple Watch dataset (Hellinger distance between red and blue curves = 0.303). On the other hand, the difference between REM and NREM was less evident in the MESA dataset (Hellinger distance = 0.187, Fig 6F). These results indicate that the dynamics underlying HR during REM and NREM sleep may be disrupted among older subjects in the MESA dataset, negatively affecting TDA’s effectiveness in differentiating REM and NREM sleep.

**Fig 6.**
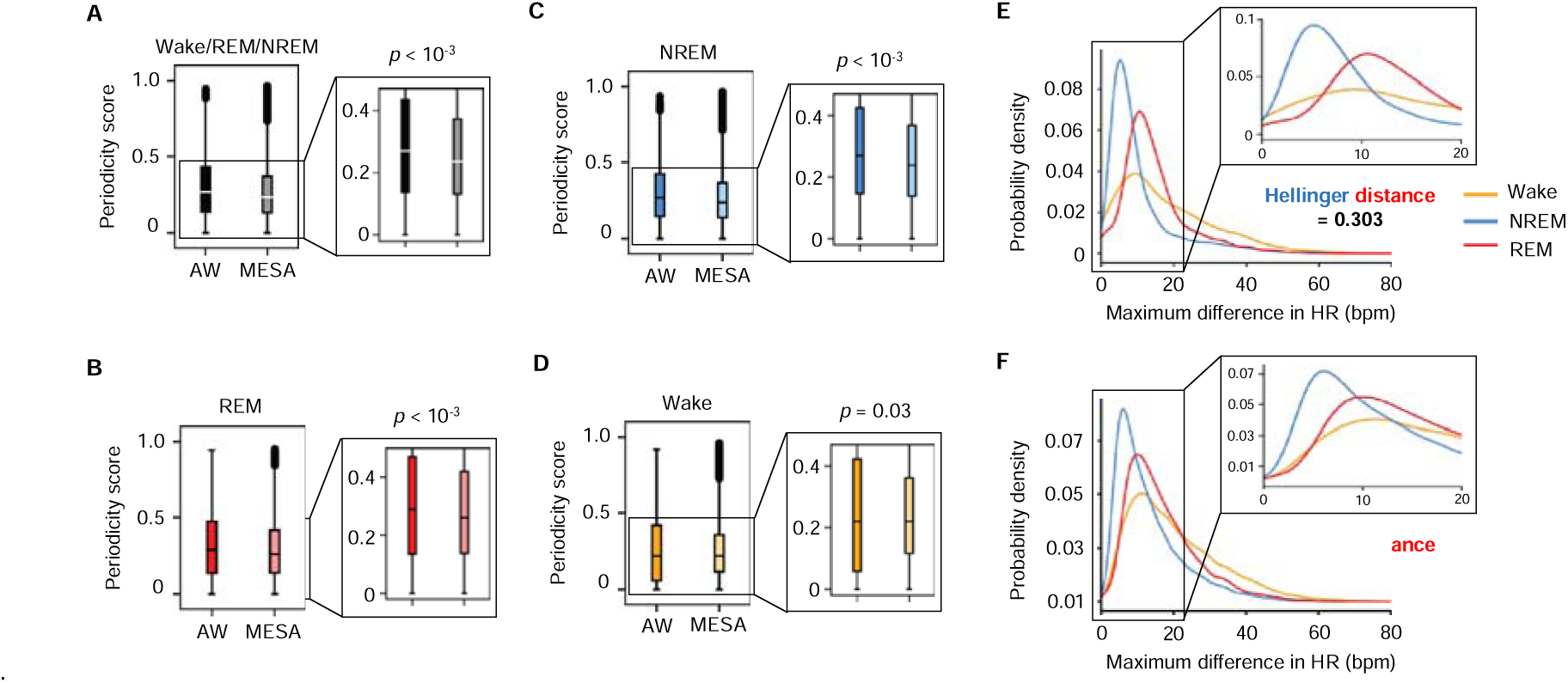
The local periodicity and the maximum differences in the HR data for each epoch. (A-D) Box-and-whisker plots of local periodicity scores for all sleep stages (A), REM (B), NREM (C), and wake (D) computed from the AW HR dataset for younger individuals and the MESA HR dataset for older individuals. Overall, the periodicity score for REM and NREM sleep was significantly higher in the AW data compared to the MESA HR data (B and C, *p*-value < 10^-3^). **(E-F)** Smooth kernel density estimate of probability distributions for maximum difference in HR for each sleep stage in the AW (E) and MESA dataset (F). The Hellinger distance between estimated probability density distributions for REM (red curve) and NREM (blue curve) sleep is significantly higher in the AW dataset compared to the MESA dataset (0.303 vs 0.187, E and F inset, respectively).

## Discussion

In this study, we proposed the use of TDA to analyze physiological signals collected from consumer-grade wearable devices. Our findings revealed a significant difference in HR TFs between REM and NREM sleep in younger healthy individuals. Thus, incorporating TFs as input features along with model-driven CPs into the neural network classifier for sleep stage prediction effectively improved Wake/REM/NREM classification performance, particularly in identifying REM and NREM sleep (Figs 2,3, and 4).

To facilitate personalized sleep medicine outside of laboratory settings, researchers have recently proposed various approaches for sleep stage prediction based on consumer-grade wearables. For instance, for the wake/REM/NREM classification, our previous algorithm based solely on the raw features and the motion CP demonstrated ∼65% balanced accuracy for REM and NREM when the accuracy of wake was fixed at 60%, with the corresponding Cohen’s K of 0.277^9^. Compared to our previous work, adding the HF CP and TFs collectively improved the balanced REM/NREM accuracy to ∼72% and corresponding Cohen’s K value to 0.434 (Table 2). This demonstrates that our algorithm can accurately identify sleep stages with higher accuracy and Cohen’s K statistics, indicating that predictions from our method are less likely to agree with the ground truth PSG recordings by chance. Importantly, our algorithm’s performance based on wearable data now closely matches the result from a previously developed sleep stage classification algorithm based on ECG data (overall accuracy = 73%, Cohen’s K = 0.460)^65^.

The key component in our algorithm’s effectiveness was the collective use of TDA and mathematical modeling. To the best of our knowledge, our work is the first study to consider applying TDA to extract hidden information embedded in the wearable data for sleep stage prediction. Moreover, using TDA, we also found the aging effects on the dynamics underlying HR for REM and NREM sleep. This may be supported by previous experimental studies reporting that older individuals have overall reduced spectral amplitude in HR^66^ or more reduced complexity of cardiovascular control during REM sleep compared to younger individuals^67,68^. Our study illustrates the effectiveness and versatility of TDA, suggesting that it can open a new avenue for wearable data science: given the robust nature of topological methods against noise, TDA can be particularly effective in analyzing highly noisy wearable sensor data and uncovering insights previously unobservable with traditional methods.

Our previous work^9^ and other studies^69^ demonstrated that the motion CP representing the changing drives in the core body temperature can be effective in sleep stages classification. In addition to the motion CP, our study further leveraged the HR CP, which was estimated using a recently proposed Bayesian MCMC-based method^42^, and found the potential of the HR CP in sleep stage prediction. This suggests that using both the two CPs, along with the TFs, may lead to improved accuracy in capturing circadian dynamics and homeostatic sleep pressure, resulting in better classification results.

Despite the strengths of our study, there exist limitations. For instance, our algorithm currently uses the MCMC framework^42^ to extract the HR CP, which is computationally demanding. This difficulty can be mitigated by replacing the MCMC framework with a recently developed non-linear least squares-based method for analyzing real-world heart rate dynamics from wearable data^45^. Moreover, despite the effectiveness of TFs in the Apple Watch dataset, its lack of effectiveness in the MESA dataset indicates that our algorithm based on the persistent diagram may not have universal applicability, especially when dealing with older individuals. Thus, further validation studies using more sophisticated tools of TDA, such as the persistence landscape^70^ or persistence image^71,72^, would be necessary.

## Simulation

All the simulations were performed using Python and MATLAB on a computing cluster, the Great Lakes Slurm cluster, which is part of the University of Michigan’s computing infrastructure, accessible at: https://arc.umich.edu/greatlakes/.

## Data Availability

The Apple Watch dataset is available on PhysioNet: https://physionet.org/content/sleep-accel/1.0.0/. The MESA dataset is available on the data repository of National Sleep Research Resource: https://sleepdata.org/datasets/mesa/files/polysomnography.

## Code Availability

The Python codes used in this study are available in the following dataset: The GitHub link will be provided here once the study is accepted.

## Declaration of conflicts of interest

O.W. and D.B.F. serve as the CEO and CSO of Arcascope, a mobile software development company for circadian rhythms. The other authors declare that they have no conflict of interest.

## Acknowledgments

This work was funded by Human Frontiers Science Program Organization grant no. RGP0019/2018, NSF DMS grant no. 2052499 and ARO MURI grant no. W911NF-22-1-0223.

## Author Contribution

M.P.L., D.W.K., and D.B.F. designed the study. M.P.L. performed and D.W.K contributed to computational modeling and simulation. All authors analyzed the data. M.P.L. and D.W.K. wrote the manuscript, and all the authors contributed to reviewing the manuscript.

