## Supplementary materials for "The combination of topological data analysis and mathematical modeling improves sleep stage prediction from consumer-grade wearables"

[†] *These authors contributed equally.*

### 1 Details regarding the Apple Watch dataset

39 Participants were recruited by the University of Michigan Sleep and Chronophysiology laboratory following the approval from the University of Michigan Institutional Review Board. Questionnaires were given to the recruited participants at the beginning of the study, and participants who have sleep-related, neurological, or psychiatric impairments that could significantly affect data collection were excluded from the study. Participants who worked on a night shift, traveled greater than two time zones within a month prior to the beginning of the ambulatory data collection period, or experienced excessive daytime sleepiness as determined by the Epworth Sleepiness Scale were further excluded from the study. The remaining subjects were instructed to wear a wrist-worn Apple Watch (Series 2 and 3, Apple Inc.) throughout the 7 to 14 days of the ambulatory recording period, except in unavoidable circumstances such as battery charging or showering. During the ambulatory recording period, Apple Watch collected heart rate (HR) data (in units of beats per minute (bpm)) using photoplethysmography (PPG) and triaxial motion data in $x$, $y$, and $z$ directions (in units of g = 9.8 $m/s^2$) using microelectromechanical systems (MEMS) type accelerometer on the dorsal side of the wrist.

On the final night of the ambulatory recording period, participants showed up at the University of Michigan Sleep and Chronophysiology Laboratory and underwent PSG recording in the presence of a registered polysomnographic technologist. During PSG recording, participants were instructed to simultaneously wear an Apple Watch, which transmitted the HR and motion data in real-time to a server at the University of Michigan. After PSG recording, participants whose PSG findings were indicative of possible Rapid Eye Movement (REM) sleep disorder or Obstructive Sleep Apnea (OSA) were identified by a board-certified sleep medicine physician and removed from all analyses. PSG recording and Apple Watch HR and motion data from all remaining 31 participants (mean age = $29.4 \pm 8.5$ years, range 19-55) were finally included in the analysis.

### 2 Mathematical formulation of persistent homology

In this section, we provide a mathematical formulation behind persistent homology. With persistent homology, we can encode, quantify, and utilize the idea of $p$-dimensional holes hidden inside the data. To do so, an algebraic representation of "holes", i.e., a homology group, is needed [2, 4, 5, 7, 9]. The key steps in extracting topological features using persistent homology can be summarized as follows:

1. Point cloud data $\mathcal{X}$ in $\mathbb{R}^n$ is used as an input.

2. For each data point $v_i \in \mathcal{X}$, construct $B_\epsilon(v_i)$, a ball of radius $\epsilon$ centered at each $v_i$, where $\epsilon > 0$.

3. For each $\epsilon$, construct a simplicial complex.

4. Increase the value of $\epsilon$ and construct a filtration of simplicial complexes.

5. For each hole in the simplicial complex, record the birth radius $\epsilon_{\text{birth}}$ and death radius $\epsilon_{\text{death}}$.

6. Plot all $(\epsilon_{\text{birth}}, \epsilon_{\text{death}})$ coordinates on a persistent diagram.

7. Using the persistent diagram, extract topological features.

We begin by introducing a simplicial complex, from which we can build the notion of chain, boundary, cycle, and homology group.

### 2.1 Simplicial complex

We first introduce simplicial complex, which is an essential tool that allows us to triangulate the given point cloud data and obtain a reasonable approximation of the underlying topological structure. Let $\mathcal{K}$ be a set of points, lines, triangles, and $p$-dimensional counterparts, where $p \in \mathbb{Z}_{\geq 0}$. We call such $p$-dimensional counterparts a $p$-simplex using the following definition.

**Definition 2.1** ($p$-simplex)**.** Let $\{v_i\}_{i \in I}$ be a collection of points in $\mathbb{R}^n$. We call each $v_i$ a vertex. Then $p$-simplex

$$\sigma_p = \langle v_0, v_1, \cdots, v_p \rangle \tag{S1}$$

is a convex hull generated by its vertices.

We call a collection of $p$-simplicies $\mathcal{K}$, a simplicial complex if it satisfies the following properties.

**Definition 2.2** (Simplicial complex)**.** Let $\mathcal{S} \subseteq \mathcal{K}$ and $\sigma \in \mathcal{K}$ be an arbitrary $p$-simplex in $\mathcal{K}$. Then $\mathcal{K}$ is a simplicial complex if

1. For all $\sigma \in \mathcal{S}$ and $\tau \subseteq \sigma$, we have $\tau \in \mathcal{K}$

2. For all $\sigma_1, \sigma_2 \in \mathcal{K}$, we have $\sigma_1 \cap \sigma_2 \in \mathcal{K}$.

In other words, $\mathcal{K}$ is a simplicial complex if (i) for any given $p$-simplex in $\mathcal{K}$, an arbitrary face of such $p$-simplex must also be a $p$-simplex in $\mathcal{K}$ and (ii) an intersection of any two simplices must also be a valid simplex in $\mathcal{K}$. There are two common methods for constructing a simplicial complex: *Vietoris-Rips Complex* and *Čech Complex*. First, the Čech complex is a generalized version of the Vietoris Rips complex.

**Definition 2.3** (Čech complex)**.** Let $\sigma = \langle v_0, \cdots, v_i, \cdots v_p \rangle$ be an arbitrary $p$-simplex and $B_\epsilon(v_i)$ be a ball with radius $\epsilon$ centered at each vertex $V_i$. We define

$$\check{C}ech(\mathcal{X}, \epsilon) = \bigcup_{p \in \mathbb{Z}_{\geq 0}} \left\{ \langle v_0, \cdots, v_p \rangle \middle| \bigcap_{i \in I} B_\epsilon(v_i) \neq \phi \right\} \tag{S2}$$

a Čech complex.

We add a $p$-simplex $\sigma$ to a Čech complex if a finite intersection of all $\epsilon$-ball around each vertex that generates $\sigma$ is non-empty. However, its implementation is often computationally inefficient because every arbitrary intersection of all $\epsilon$-ball needs to be evaluated. Hence, a more computationally efficient method that still provides a reasonable approximation of the underlying topological space is needed. Vietoris-Rips complex accomplishes this.

**Definition 2.4** (Vietoris-Rips complex)**.** Given an arbitrary $p$-simplex $\sigma = \langle v_0, \cdots, v_i, \cdots, v_p \rangle$ and a metric $d : \mathbb{R}^n \times \mathbb{R}^n \to \mathbb{R}$, we define

$$Rips(\mathcal{X}, \epsilon) = \bigcup_{p \in \mathbb{Z}_{\geq 0}} \left\{ \langle v_0, \cdots, v_p \rangle \middle| d(v_i, v_j) \leq \epsilon, 0 \leq i, j \leq p \right\} \tag{S3}$$

a Vietoris Rips complex.

Note that Vietoris Rips complex only requires every *pairwise distance* between vertices to be at most $\epsilon$. Thus, the Vietoris Rips complex much more computationally efficient than the Čech complex, but the construction of Vietoris Rips is comparably weaker than that of the Čech complex, making. This makes the Vietoris Rips complex more suitable for application in Topological Data Analysis, provided that the Vietoris Rips complex gives a reasonable approximation of the topological space generated by the input data set. Therefore, we need specific criteria to determine if and when the Vietoris Rips complex is appropriate. The following theorem provides this criterion:

**Theorem 1.** *Given* $\epsilon \in \mathbb{R}^+$,

$$\check{C}ech(\epsilon) \subseteq Rips(\epsilon) \subseteq \check{C}ech(2\epsilon) \tag{S4}$$

Hence, given that Čech complex provides a good approximation of topological space generated by the data for both $\epsilon$ and $2\epsilon$, we can assert that Vietoris Rips complex for $\epsilon$ gives a reasonable approximation of the underlying topological structure. In this study, we implemented the Vietoris Rips complex using the Giotto-TDA package in Python 3.9 [8].

### 2.2 Chain, Cycle, Boundary, and Homology Group

Using $p$-dimensional complex, we now define chain, cycle, and boundary group to formalize the notion of $p$-dimensional "hole", i.e., homology group. We first define algebraic representations of simplicies by introducing $p$-chain.

**Definition 2.5** ($p$-chain)**.** Given a collection of $p$-simplex $\{\sigma_i\}_{i\in I}$ from a simplicial complex $\mathcal{K}$, a $p$-chain $C_p$ is a finite sum of $\sigma_i$s multiplied with some arbitrary coefficient in module 2 $\mathbb{Z}_2$. In other words,

$$C_p = \sum_{i\in I}^{n} c_i\sigma_i, \text{ where } c_i \in \mathbb{Z}_2, \sigma_i \in \mathcal{K}, dim(\sigma_i) = p \tag{S5}$$

A $p$-chain group is simply a collection of all $p$-chains in a given simplicial complex.

**Definition 2.6** ($p$-chain group)**.** Given a simplicial complex $\mathcal{K}$ and $\sigma_i \in \mathcal{K}$,

$$C_p(\mathcal{K}) = \left\{ \sum_{i\in I}^{n} c_i\sigma_i \middle| c_i \in \mathbb{Z}_2, dim(\sigma_i) = p \right\} \tag{S6}$$

is a $p$-chain group.

Note that $c_i$'s are coefficients from modulo 2 coset $\mathbb{Z}_2$. Hence, for any arbitrary simplex $\sigma \in \mathcal{K}$, $\sigma + \sigma = 0$. For example, consider two $p$-chains $C_p = \sigma_1 + \sigma_2 + \sigma_3$ and $C_p^{'} = \sigma_1 - \sigma_2 - \sigma_4$, where each $\sigma_{1,\cdots,4}$ is a $p$-simplex in a given simplicial complex $\mathcal{K}$. Then $C_p + C_p^{'} = (\sigma_1 - \sigma_2 + \sigma_3) + (\sigma_1 - \sigma_2 - \sigma_4) = \sigma_3 - \sigma_4$.

Now we consider the cycle group and boundary group. To do so, we first define the boundary operator $\partial_p$, which provides a nested relationship between a sequence of chain groups.

**Definition 2.7** (Boundary operator)**.** Given a simplicial complex $\mathcal{K}$ and $p$-simplex $\sigma = \langle v_0, v_1, \cdots, v_p\rangle$, the boundary operator $\partial_p : C_p(\mathcal{K}) \to C_{p-1}(\mathcal{K})$ is a group homomorphism that assigns $\sigma$ to its boundary:

$$\partial_p(\sigma) = \partial_p(\langle v_0, \cdots, v_p\rangle) = \langle v_0, \cdots, \hat{v_i}, \cdots, v_p\rangle \tag{S7}$$

Note that $\hat{\cdot}$ is a dropout operator, so for some $0 \leq i \leq p$, the vertex $v_i$ is droped out under $\partial_p$, which indicates that $\langle v_0, \cdots, \hat{v_i}, \cdots, v_p\rangle \in C_{p-1}(\mathcal{K})$. The boundary operator $\partial_p$ gives a nested relation among chain groups $C_p(\mathcal{K})$ for each $p$. This nested relation among all chain groups is called a *chain complex*:

$$\cdots C_n(\mathcal{K}) \xrightarrow{\partial_n} C_{n-1}(\mathcal{K}) \xrightarrow{\partial_{n-1}} \cdots \xrightarrow{\partial_2} C_1(\mathcal{K}) \xrightarrow{\partial_1} C_0(\mathcal{K}) \xrightarrow{\partial_0} 0 \tag{S8}$$

One fundamental result of chain complex is that the image of the boundary does not have a boundary.

**Theorem 2** (Munkres (1993) Lemma 5.3)**.** *For any given simplicial complex* $\mathcal{K}$ *and any* $p$*-chain* $c \in C_p(\mathcal{K})$*, we have*

$$(\partial_{p-1} \circ \partial_p)(c) = 0 \tag{S9}$$

*for all* $p \in \mathbb{Z}_{\geq 0}$ *[6].*

This fundamental result motivates the definition of boundary group $B_p(\mathcal{K})$.

**Definition 2.8** ($p$-boundary)**.** Given chain groups $C_p, C_{p+1}$ and a boundary operator $\partial_{p+1} : C_{p+1} \to C_p$ of a simplicial complex $\mathcal{K}$, a $p$-chain $\sigma_p$ such that

$$\sigma_p = \langle v_0, \cdots, \hat{v_i}, \cdots, v_{p+1} \rangle = \partial_{p+1}(\sigma_{p+1}) \tag{S10}$$

is a $p$-boundary.

We note that a $p$-simplex $\sigma_p$ is a $p$-boundary if and only if it is an image of some $(p+1)$-chain under $(p+1)$-boundary operator $\partial_{p+1}$.

**Definition 2.9** ($p$-boundary group)**.** Given a simplicial complex $\mathcal{K}$,

$$B_p(\mathcal{K}) := \Big\{ \sigma_p \Big| \sigma_p \in im(\partial_{p+1}) \Big\} \tag{S11}$$

is a $p$-boundary group. Hence,

$$B_p(\mathcal{K}) = im(\partial_{p+1}) \tag{S12}$$

A $p$-cycle is a $p$-simplex in a simplicial simplex such that its image under the $p$-boundary operator is a zero, i.e, its boundary is a zero.

**Definition 2.10** ($p$-cycle)**.** Given a simplicial complex $\mathcal{K}$, a $p$-cycle is a $p$-simplex $\sigma_p \in C_p(\mathcal{K})$ such that $\partial_p(\sigma_p) = 0$.

Thus, each $p$-cycle is in the kernel of a $p$-boundary operator. Hence, a $p$-cycle group is equivalent to the kernel of $p$-boundary operator.

**Definition 2.11** ($p$-cycle group)**.** A $p$-cycle group $B_p(\mathcal{K})$ is a collection of all $p$-cycles, i.e,

$$Z_p(\mathcal{K}) := \Big\{ \sigma_p \in C_p(\mathcal{K}) \Big| \partial_p(\sigma_p) = 0 \Big\} = ker(\partial_p) \tag{S13}$$

Defining the chain, cycle, and boundary group now allows us to introduce the homology group, which formalizes the notion of holes. We first note that the chain complex of boundary operators, combined with Theorem 2 from [6], suggest that for any $p \in \mathbb{Z}_{\geq 0}, \partial_p \circ \partial_{p+1} = 0$. This implies that for any $\sigma_p \in B_p(\mathcal{K}) = im(\partial_{p+1})$, we have $(\partial_p \circ \partial_{p+1})(\sigma_p) = 0$ as $\sigma_p$ is a boundary of some $(p+1)$-simplex $\sigma_{p+1}$. Therefore,

$$im(\partial_{p+1}) \subseteq ker(\partial_p) \tag{S14}$$

which is equivalent to

$$B_p(\mathcal{K}) \subseteq Z_p(\mathcal{K}) \tag{S15}$$

This result finally allows us to define $p$-homology group.

**Definition 2.12** ($p$-homology Group)**.** Given a $p$-boundary group $B_p(\mathcal{K})$ and a $p$-cycle group $Z_p(\mathcal{K})$, a $p$-homology group is a quotient space obtained by identifying all $p$-boundaries together in $Z_p(\mathcal{K})$. In other words,

$$H_p(\mathcal{K}) := Z_p(\mathcal{K})/B_p(\mathcal{K}) = ker(\partial_p)/im(\partial_{p+1}) \tag{S16}$$

A $p$-homology group for a given simplicial complex $\mathcal{K}$ formalizes the notion of $p$-dimensional hole. We can also quantify the number of different $p$-dimensional holes in $\mathcal{K}$ with $p$-dimensional Betti numbers.

**Definition 2.13** ($p$-Betti number)**.** Given a $p$-homology group $H_p(\mathcal{K})$, Betti number is a dimension of $p$-homology group:

$$\beta_p(\mathcal{K}) = rank(H_p(\mathcal{K})) \tag{S17}$$

Informally, $\beta_0(\mathcal{K})$ denotes the number of connected components generated in the simplicial complex; $\beta_1(\mathcal{K})$ denotes the number of loops; and $\beta_2(\mathcal{K})$ denotes the number of voids, and so on. In many practical application of persistent homology, $H_p(\mathcal{K})$ and $\beta_p(\mathcal{K})$ for $p \geq 3$ are rarely considered, mainly due to computational inefficiency of computing $H_p$ and $\beta_p$ for $p \geq 3$.

### 2.3 Persistent homology

Persistent homology generalizes the notion of a homology group to a nested sequence of simplicial complexes and tracks the timing of birth and death of each hole. Hence, we formalize the idea of *birth* and *death* of each homology group. First, we need a nested relation of simplicial complexes called filtration.

**Definition 2.14** (Filtration)**.** Let $\{\mathcal{K}_i\}_{i\in I}$ be a collection of simplicial complexes. Then the nested relation among simplicial complexes satisfying

$$\phi = \mathcal{K}_0 \subseteq \mathcal{K}_1 \subseteq \cdots \subseteq \mathcal{K}_i \subseteq \cdots \subseteq \mathcal{K}_j \subseteq \cdots \subseteq \mathcal{K}_n \tag{S18}$$

for all $i \leq j$ is called a filtration of simplicial complexes.

Intuitively, one can think of adding a collection of $p$-simplicies to an existing simplicial complex $\mathcal{K}_i$ to construct the subsequent simplicial complex $\mathcal{K}_{i+1}$ in a filtration, and so on. The core motivation behind persistent homology is to observe the evolution of $p$-dimensional homology group and betti number in a filtration of simplicial complexes. Hence, a mapping between simplicial complexes in filtration is needed:

**Definition 2.15** (Inclusion map)**.** For each $i \in I$, let

$$\hookrightarrow_i \ : \mathcal{K}_i \to \mathcal{K}_{i+1} \tag{S19}$$

be the inclusion map from $\mathcal{K}_i$ to its subsequent simplicial complex $\mathcal{K}_{i+1}$ within a filtration.

The inclusion map $\hookrightarrow_i$ thus induces a homomorphism $f_p^{i,j} : H_p(\mathcal{K}_i) \to H_p(\mathcal{K}_j)$ where

$$f_p^{i,j} = \ \hookrightarrow_{j-1} \circ \cdots \circ \hookrightarrow_{i+1} \circ \hookrightarrow_i \tag{S20}$$

between homology group of any two simplicial complexes $\mathcal{K}_i$ and $\mathcal{K}_j$, for each dimension $p$. This allows us to define $p$-dimensional persistent homology, which will then formalize the *birth* and *death* events of holes in a filtration.

**Definition 2.16** ($p$-dimensional persistent homology)**.** Let $f_p^{i,j} : H_p(\mathcal{K}_i) \to H_p(\mathcal{K}_j)$ be a group homomorphism between $p$-homology group of simplicial complexes $\mathcal{K}_i$ and $\mathcal{K}_j$. Then for $0 \leq i \leq j$, $p$-dimensional persistent homology is defined as

$$H_p^{i,j} := im(f_p^{i,j}) \tag{S21}$$

for each dimension $p \in \mathbb{Z}_{\geq 0}$.

In particular, we observe that $p$-dimensional persistent homology $H_p^{i,j}$ is a generalization of $p$-homology group, since $i = j$ implies $f_p^{i,j}$ maps from $H_p(\mathcal{K}_i)$ to itself, which therefore suggests $im(f_p^{i,j}) = im(f_p^{i,i}) = H_p(\mathcal{K}_i)$. A $p$-dimensional persistent homology defines the birth moment of $p$-dimensional holes. Specifically, if we let $\gamma_p$ be a $p$-dimensional hole, we say that $\gamma_p$ is born at $\mathcal{K}_i$ if

$$\gamma_p \in H_p(\mathcal{K}_i), \text{ but } \gamma_p \notin im(f_p^{i-1,i}) = H_p^{i-1,i} \tag{S22}$$

Recall that $f_p$ is a composition of inclusion maps between $p$ homology classes, as shown in equation (S20). Similarly, if $\gamma_p$ is born at $\mathcal{K}_i$, then it dies at $\mathcal{K}_j$ if

$$f_p^{i,j-1}(\gamma_p) \notin im(f_p^{i-1,j-1}) = H_p^{i-1,j-1}, \text{ but } f_p^{j-1,j}(\gamma_p) \in im(f_p^{j-1,j}) = H_p^{j-1,j} \tag{S23}$$

We see that $\gamma_p$ is born at $\mathcal{K}_i$ as it is contained in the homology group of $\mathcal{K}_i$, but is not contained in the image of the inclusion map from the homology group of $\mathcal{K}_{i-1}$ to homology group of $\mathcal{K}_i$. In other words, because $\gamma_p$ is not contained in the persistence homology from $\mathcal{K}_{i-1}$ to $\mathcal{K}_i$, $\gamma_p$ was not existent before $\mathcal{K}_i$. The fact that $\gamma_p$ is existent in the homology group of $\mathcal{K}_i$ asserts that $\gamma_p$ was born during the construction of $\mathcal{K}_i$ from $\mathcal{K}_{i-1}$. Moreover, the absence of $\gamma_p$ in $H_p(\mathcal{K}_{i-1})$ implies that the image of $f_p^{i-1,k}$ does not contain $f_p^{i,k}(\gamma_p)$ for all $k \geq i$, which means that $\gamma_p$ is no longer present in the persistent homology $H_p^{i,k}$. Because $k = j$ is the first time that $f_p^{i,k}(\gamma_p) \in im(f_p^{i-1,k}) = H_p^{i-1,k}$, $\gamma_p$ dies entering $\mathcal{K}_j$. Finally, we now define the persistence of a $p$-dimensional hole.

**Definition 2.17** (Persistence of a $p$-dimensional hole)**.** Given a $p$-dimensional hole $\gamma_p$ that is born at $\mathcal{K}_i$ and dies at $\mathcal{K}_j$, the persistence of $\gamma_p$ is defined as

$$pers(\gamma_p) = j - i \tag{S24}$$

Note that if $j = +\infty$, then $pers(\gamma_p) = +\infty$, which means $\gamma_p$ never disappears in the filtration. There will always be a 0-dimensional hole (connected component) with an infinite persistence, and such a 0-dimensional hole with infinite persistence is often disregarded in the analysis.

A $p$-dimensional persistence diagram $Dgm_p(f_p)$ visualizes all $p$-dimensional persistence homology along the filtration $\{\mathcal{K}_\epsilon\}$ for all $\epsilon \in \mathbb{R}^+$ [1, 3, 9]. Each $p$-dimensional persistence diagram is a collection of points $(\epsilon_i, \epsilon_j)$ that encode the respective birth and death event for each $p$-dimensional hole in a filtration (see main text Fig 1B). Thus, each $(\epsilon_i, \epsilon_j)$ represents a $p$-dimensional hole that is born at $\mathcal{K}_{\epsilon_i}$ and dies at $\mathcal{K}_{\epsilon_j}$. The horizontal axis of a persistence diagram represents the birth epsilon values of holes, while the vertical axis represents the death epsilon values. Note that since the birth and death of all holes are defined for $i \leq j$, every $(\epsilon_i, \epsilon_j)$ are plotted above the diagonal on the persistence diagram.

### Supplementary Figures

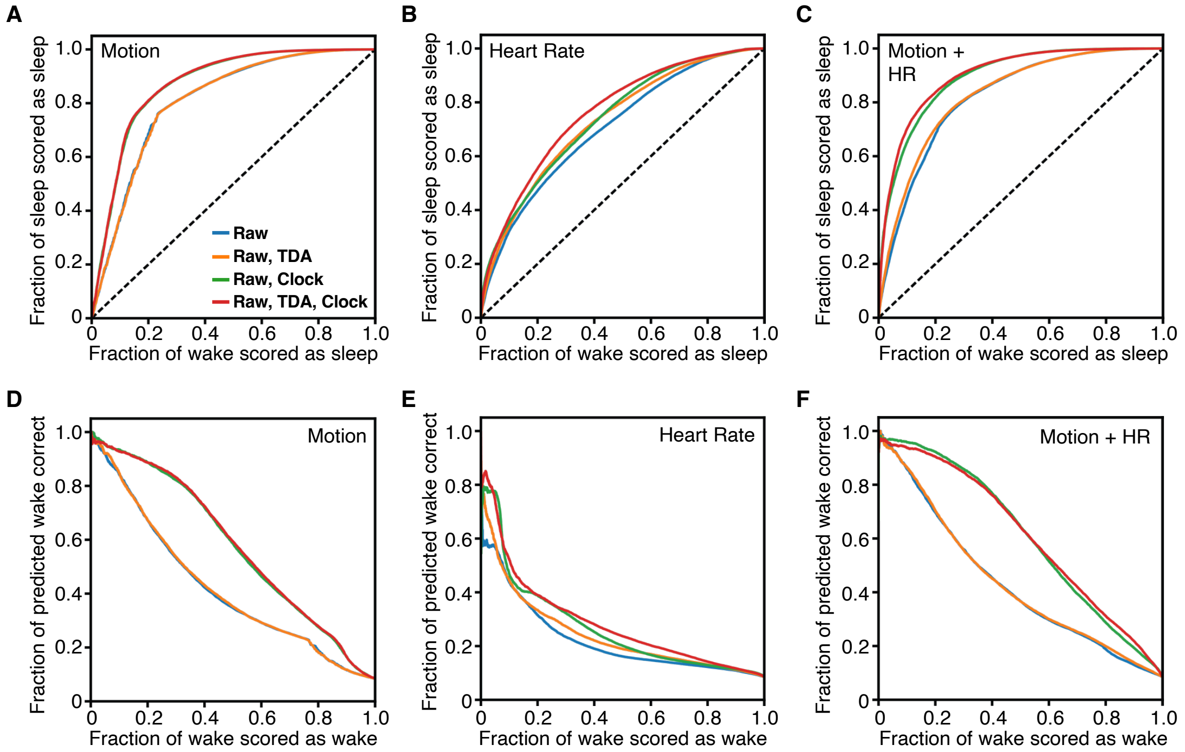


**Fig. S1**: **(A) ROC and PRC curves for sleep/wake classification within the Apple Watch Dataset. (A-C)** ROC curves for sleep/wake classification using motion-derived features (A), heart rate derived features (B) and both motion and heart rate-derived features (C). **(D-F)** PRC curves for sleep/wake classification using motion-derived features (D), heart rate derived features (E) and both motion and heart rate-derived features (F).

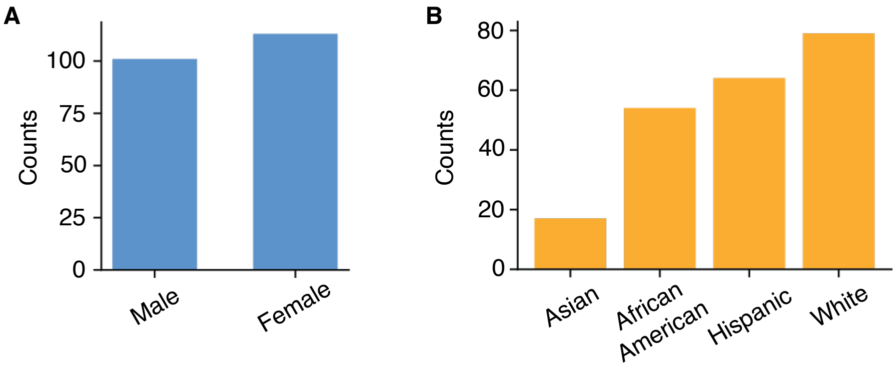


**Fig. S2**: **Demographic information for MESA dataset cohorts analyzed in this study.** **(A)** A histogram showing the sex information of the MESA subjects. **(B)** A histogram showing race information of the MESA subjects.

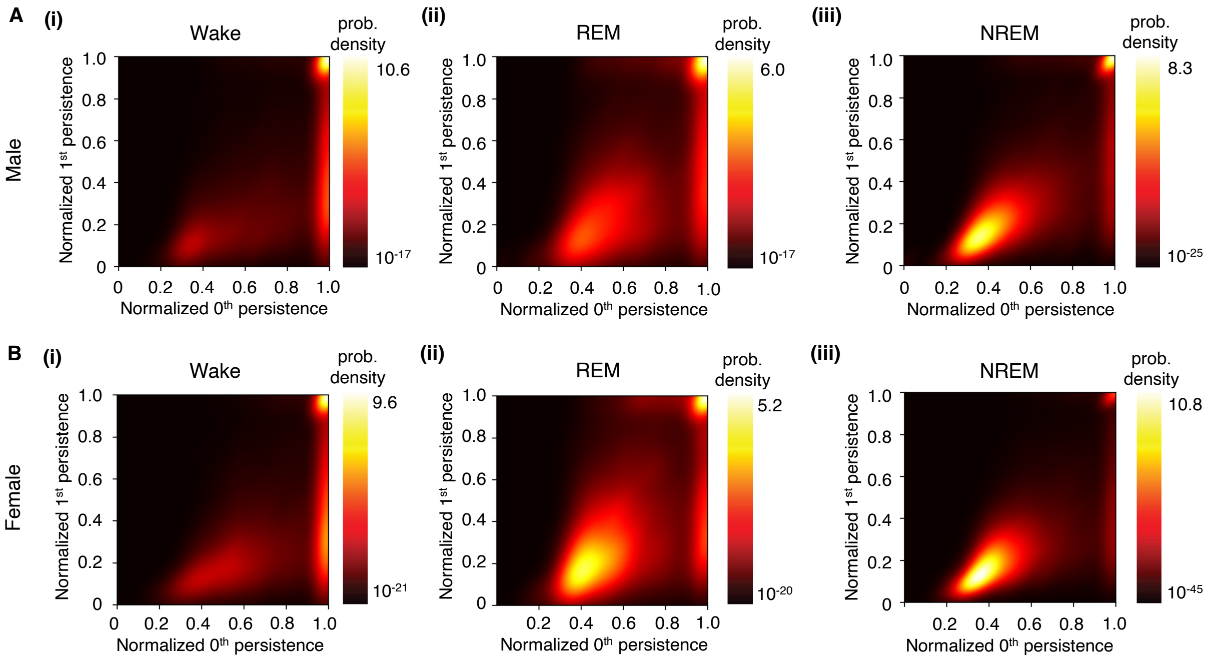


**Fig. S3**: **Probability density distributions of heart rate topological features among MESA subjects based on sex.** **(A)** Smooth kernel density estimates of the distribution of topological features for wake (i), REM (ii), and NREM (iii) in male participants. **(B)** Smooth kernel density estimates of topological features for wake (i), REM (ii), and NREM (iii) in female participants. The distribution of HR topological features is similar across different subjects for both male and female subjects.

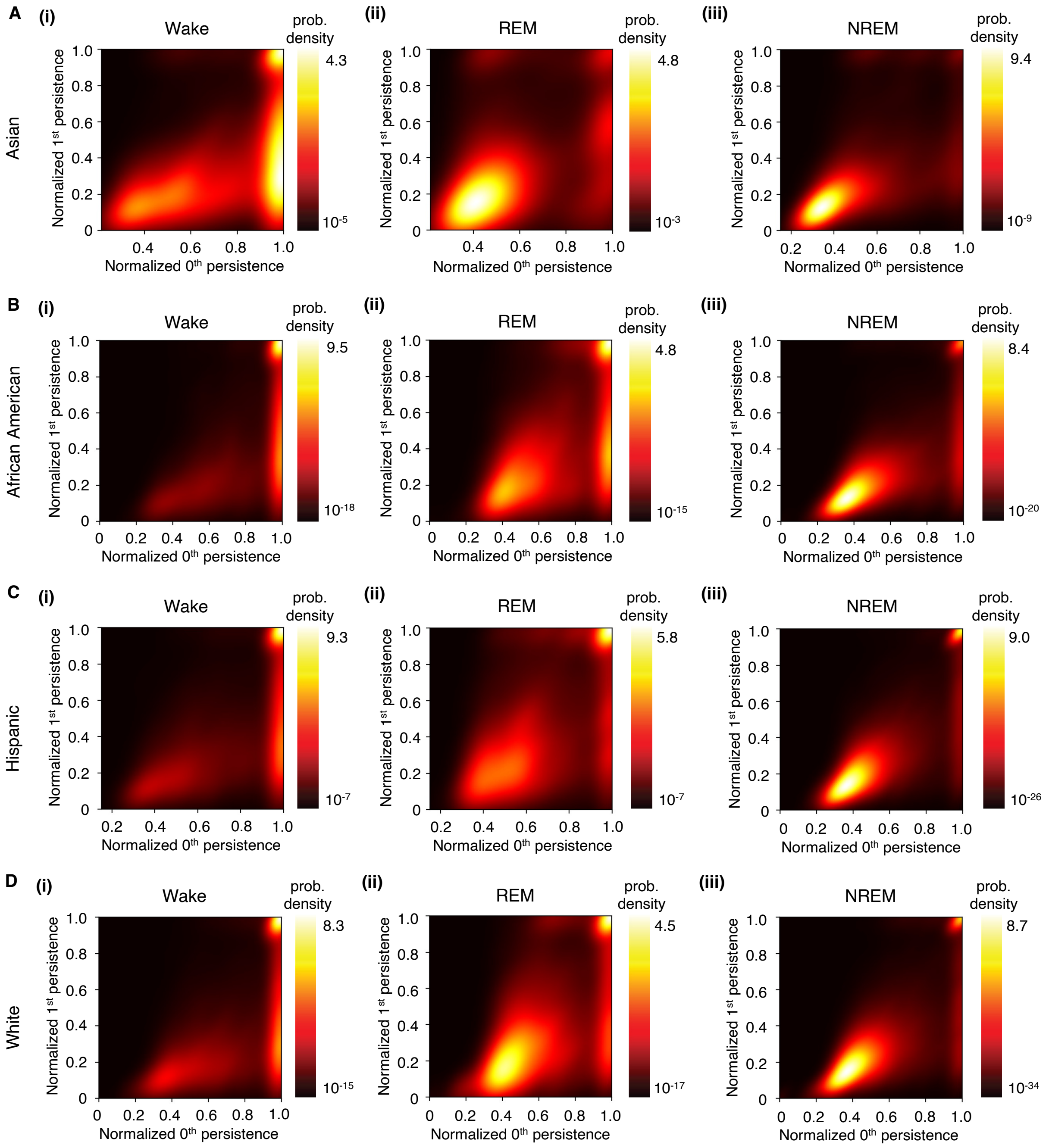


**Fig. S4**: **Probability density distributions of heart rate topological features among MESA subjects based on race.** **(A-D)** Smooth kernel density estimates of the distribution of topological features for Asian (A), African American (B), Hispanic (C), and White (D) participants during wake (i), REM (ii), and NREM (iii). The distribution of HR topological features is similar across different subjects regardless of the subject's race information.

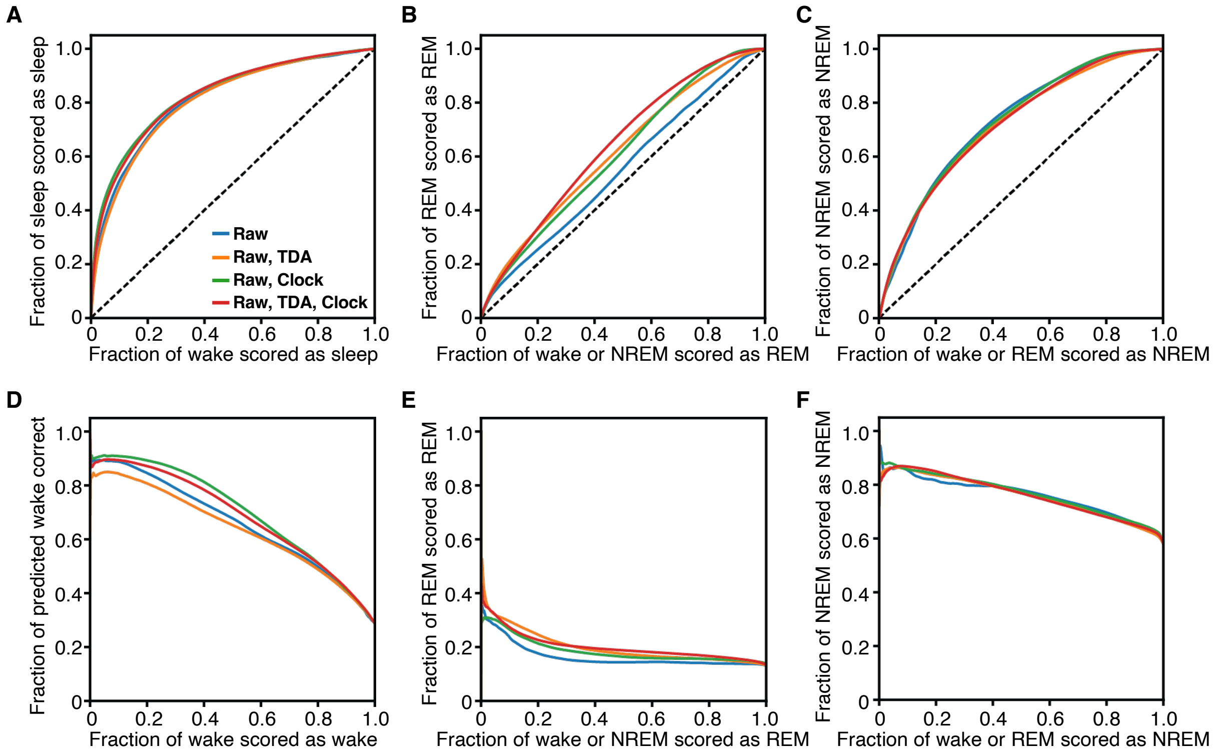


**Fig. S5**: **One-versus-rest ROC and PRC curves for wake/REM/NREM classification within the MESA subjects. (A-C)** One-versus-rest ROC curves for wake/REM/NREM classification for wake (A), REM (B) and NREM (C) using different features within the MESA dataset. **(D-F)** One-versus-rest PRC curves for wake/REM/NREM for wake (A), REM (B) and NREM (C) using different features within the MESA dataset. Unlike in the Apple Watch dataset, topological features failed to significantly improve classification performance across different stages for the MESA dataset.

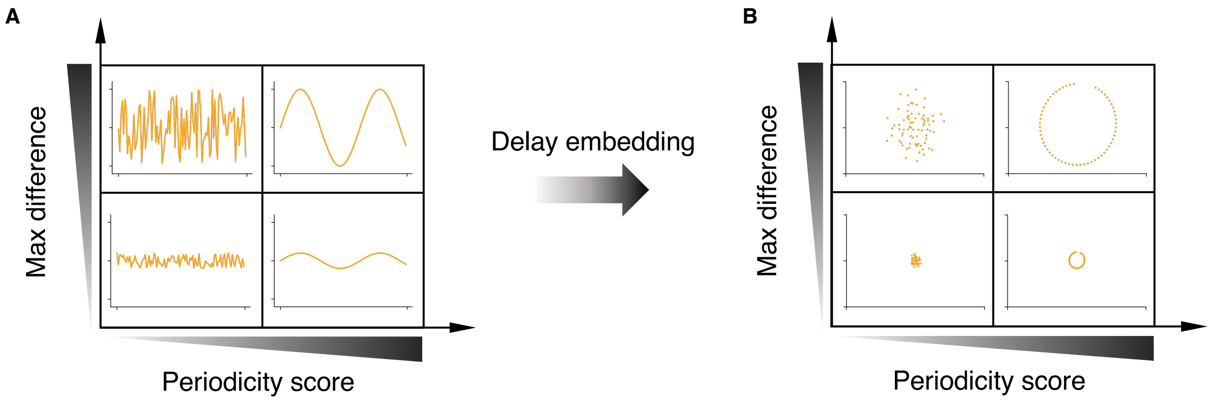


**Fig. S6**: **Relationship between the size and circularity of high dimensional hole to the periodicity score and maximum difference in time-series. (A)** Example of time-series with different values of periodicity score and maximum difference in time-series data. **(B)** After delay embedding using optimal embedding parameters, a high dimensional hole represented by the 1st and 2nd principal components ($x$ and $y$ axis, respectively), can be better reconstructed when the periodicity and a maximum difference of the time-series are high.

**Table S1**: **Hyperparameters search table.** The reported hyperparameters for the neural network model were found by optimizing the overall AUROC and AUPRC values. The same hyperparameters were used for all simulations in all datasets.

| **Hyperparameters** | |
|---|---|
| Solver | ADAM |
| Activation function | arctan |
| Output function | Sigmoid |
| Hidden layer dimension | (50,50,50) |
| $L_2$ regularization | 0.001 |
| Max number of iterations | 500 |
| Early stopping | True |

**Table S2**: **Sleep/wake sleep prediction results within the Apple Watch dataset.** The reported AUROC and AUPRC values were calculated from the ROC and PRC curves for sleep/wake differentiation within the Apple Watch dataset using the MCCV (Supplementary Fig S1).

| | Types of features used | | | |
|---|---|---|---|---|
| | Raw | Raw, TDA | Raw, clock | Raw, TDA, clock |
| **Motion only** | | | | |
| AUROC | 0.810 | 0.809 | 0.871 | 0.872 |
| AUPRC | 0.420 | 0.422 | 0.576 | 0.580 |
| **HR only** | | | | |
| AUROC | 0.706 | 0.732 | 0.739 | 0.763 |
| AUPRC | 0.222 | 0.243 | 0.268 | 0.287 |
| **Motion + HR** | | | | |
| AUROC | 0.820 | 0.828 | 0.893 | 0.901 |
| AUPRC | 0.434 | 0.437 | 0.609 | 0.611 |

[4] Herbert Edelsbrunner and John Harer. *Computational topology: an introduction.* American Mathematical Soc., 2010.

[5] Charles Epstein, Gunnar Carlsson, and Herbert Edelsbrunner. Topological data analysis. *Inverse Problems*, 27(12):120201, 2011.

[6] James R Munkres. *Elements of algebraic topology.* CRC press, 2018.

[7] Yara Skaf and Reinhard Laubenbacher. Topological data analysis in biomedicine: A review. *Journal of Biomedical Informatics*, 130:104082, 2022.

[8] Guillaume Tauzin, Umberto Lupo, Lewis Tunstall, Julian Burella Pérez, Matteo Caorsi, Anibal M Medina-Mardones, Alberto Dassatti, and Kathryn Hess. giotto-tda: A topological data analysis toolkit for machine learning and data exploration. *The Journal of Machine Learning Research*, 22(1):1834–1839, 2021.

[9] Larry Wasserman. Topological data analysis. *Annual Review of Statistics and Its Application*, 5:501–532, 2018.
